# Deep-tissue absolute force spectroscopy with sub-piconewton precision

**DOI:** 10.64898/2026.03.23.712846

**Authors:** Tatiana Merle, Amsha Proag, Ronan Bouzignac, Vanessa Dougados, Nawel Fellouah Ould Moussa, Anne Sentenac, Anne Pellissier Monier, Magali Suzanne, Thomas Mangeat

## Abstract

Quantitative measurements performed directly in vivo are necessary to understand how forces shape living tissues, yet this remains challenging due to optical scattering and mechanical complexity. Here, we present a method for making absolute force measurements using nanoscopic optical tweezers with a sensitivity of 300 fN in optically turbid biological media. Our approach combines back focal plane interferometry operating within the optical memory effect regime with a global fluctuation-dissipation fitting framework that simultaneously calibrates position detection, trap stiffness, and viscoelastic response. This method overcomes aberration-induced biases by jointly fitting passive fluctuations and driven harmonic responses, enabling robust force reconstruction in thick, scattering tissues within the mechanically relevant frequency range below 300 Hz. We validate our approach using highly scattering *Drosophila* pupae and embryos, demonstrating reliable in vivo measurements of forces and mechanical properties. Operating at a 1 kHz acquisition bandwidth, the system captures relevant mechanical dynamics without requiring extended high-frequency detection. Using this framework, we quantify the increase in cortical tension during pupal morphogenesis, characterize tissue viscoelasticity, and reveal stage-dependent variations in nuclear membrane tension during embryogenesis, even in the presence of strong ATP-driven fluctuations. Beyond bulk measurements, our method enables the quantitative mechanical characterization of single cells within mechanically coupled tissues.

## 1 Introduction

### Optical tweezers as quantitative force tools in biology

Since the invention of the gradient force optical trap by A. Ashkin [1], single-objective optical tweezers have enabled major advances in the understanding of molecular machinery, both *in vitro* and *in vivo* [2]. Optical tweezers have been widely applied in three main classes of biological studies: single-molecule manipulation *in vitro* [3, 4], multibeam holographic optical tweezers for micromanipulation [5, 6, 7], and quantitative force spectroscopy in living cells [8, 9, 10, 11]. Among these, force spectroscopy in living systems holds particular promise for mechanobiology, but also presents significant implementation and methodological challenges, especially when extended to living tissues.

### Instrumental requirements and detection strategies

Any optical tweezers setup relies on two essential optical blocks. The first block is dedicated to the generation and precise positioning of the optical trap(s) in the object plane. The second block tracks the position of the trapped object relative to the centre of the focusing beam. A wide variety of experimental implementations exists for each of these two functions, and the required level of precision strongly depends on the biological question and experimental context. Tracking the trapped object is a critical point. It can be achieved either using camera-based detection or interferometric techniques such as back-focal-plane interferometry (BFPi) [12]. Camera-based methods are well adapted to multibeam optical tweezers and holographic configurations [13], but they are limited in temporal resolution, which restricts quantitative force spectroscopy and generally prevents simultaneous high-speed fluorescence imaging. By contrast, BFPi enables object tracking at very high frequencies, up to the MHz range, and is therefore compatible with force spectroscopy. Importantly, it is currently the only method capable of distinguishing true optical trapping from indirect mechanical coupling. Moreover, BFPi provides three-dimensional tracking of the trapped object, allowing detection of tissue drift or loss of confinement and the exclusion of unreliable measurements.Recently, Catala-Castro et al. introduced Time-Shared Optical Microrheology (TimSOM)[14], which uses a single laser that is alternated between two traps to generate quasi-simultaneous stress–strain measurements using a high-frequency acousto-optic deflector (AOD). While this approach reduces alignment complexity and enforces trap symmetry, temporal multiplexing intrinsically alters probe trajectories in viscoelastic materials and requires dedicated modelling to retrieve artefact-free responses.

### Challenges specific to living tissues and trapped objects

For applications in living tissues, a first major challenge is the ability to optically trap refractive objects while limiting optical power to avoid excessive heating. Two main strategies are commonly used: the introduction of exogenous silica particles or the direct trapping of endogenous organelles with high refractive index, such as lipid droplets, cell membranes or Golgi structures. Among these options, trapping lipid droplets is particularly elegant and accessible across a wide range of living organisms. Lipid droplets are spherical organelles with a high refractive index (1.42–1.50) and sizes ranging from 100 nm to more than 1 µm. Their physical properties are sufficiently stable over the timescales of optical tweezers measurements, typically a few seconds. In addition, their surface tension is high enough to be comparable to certain nuclear or plasma membrane tensions, with ratios on the order of 100 [15].However, lipid droplets are far from monodisperse in size, exhibit variable refractive indices, and are embedded in complex viscoelastic environments in living tissues. As a consequence, optical trap calibration must be performed for each individual measurement, raising the central question of how to reliably calibrate optical tweezers under such experimentally uncontrolled conditions.

### Force calibration strategies in complex biological media

One approach to trap calibration is the direct measurement of optical momentum by quantifying the injected and diffracted components of light [16]. This strategy has been widely used in single-molecule manipulation with dual-objective 4*π* microscope systems, where nearly complete angular collection of scattered light enables accurate force estimation. However, such configurations come at the cost of high system complexity and are generally incompatible with thick biological samples, which require physical accessibility for environmental control, microfluidic integration or dedicated chambers. Single-objective implementations using adapted back-focal-plane interferometry have therefore been explored [17], but these approaches remain largely limited to thin biological samples, as thick tissues strongly scatter light and degrade the detection signal. An alternative strategy is provided by fluctuation–dissipation-based optical tweezers (FDT optical tweezers), which combine passive and laser-driven microrheology with BFPi tracking. In this framework, the fluctuation–dissipation theorem is used to simultaneously calibrate the optical trap and extract the viscoelastic properties of the surrounding medium [18, 19]. Several studies have demonstrated the robustness and reproducibility of this approach in laser-driven configurations at high frequencies. In particular, at frequencies up to several hundred hertz, the fluctuation–dissipation theorem remains valid for trapped objects in the cytoplasm. Multiplexing several driving frequencies allows a significant increase in the precision of trap stiffness calibration. Recent studies have further shown that, thanks to the combination of high-frequency laser driving and the large bandwidth of BFPi detection, it is possible to calibrate the position detector, the trap stiffness and the viscoelastic properties of the medium simultaneously within only a few seconds[20]. This directly challenges the common assumption that FDT-based approaches are intrinsically slow, an argument often raised in favour of momentum-based force detection methods.

### Scope and contribution of the present work

In this study, we propose extending BFPi-based optical tweezers to millimetre-thick biological tissues using square wave FDT calibration, whilst ensuring compatibility with high-resolution fluorescence imaging. First, we show that BFPi tracking is still possible in the presence of strong optical scattering. We then show that square wave multiplex calibration can be reliably applied in casein emulsion turbid media, using milk. Finally, we present force measurements in living tissue with sub-piconewton precision, a feat that had not previously been achieved in this context. Beyond methodological advances, a central challenge in tissue mechanobiology is extracting intrinsic single-cell mechanical properties in mechanically coupled environments. In living tissues, individual cells are mechanically connected through junctions and cytoskeleton networks. This means that local force measurements are inherently influenced by long-range force propagation from neighbouring cells. Consequently, mechanical measurements performed on a single cell cannot be interpreted as purely local responses. Here, we address this issue by combining localised optical tweezers-based microrheology with harmonic filtering. This identifies a frequency window in which the mechanical response becomes effectively localised, enabling intrinsic single-cell mechanical properties to be extracted without the need to isolate the probed cell mechanically.

## 2 10-nm precision tracking with back focal plane interferometry (BFPi) through turbid media

Back focal plane interferometry is an essential method for tracking the trapped object with reference to the center of the optical trap at nanometric scale (Fig. 1A). Theoretical and experimental studies [12, 21, 22] have shown that intensity changes in the Back Focal Plane of the objective (BFP) are a consequence of the interference between the incident focused beam and the field scattered by the trapped lipid droplet, which occurs throughout the angular range of the focus beam.

**Figure 1:**
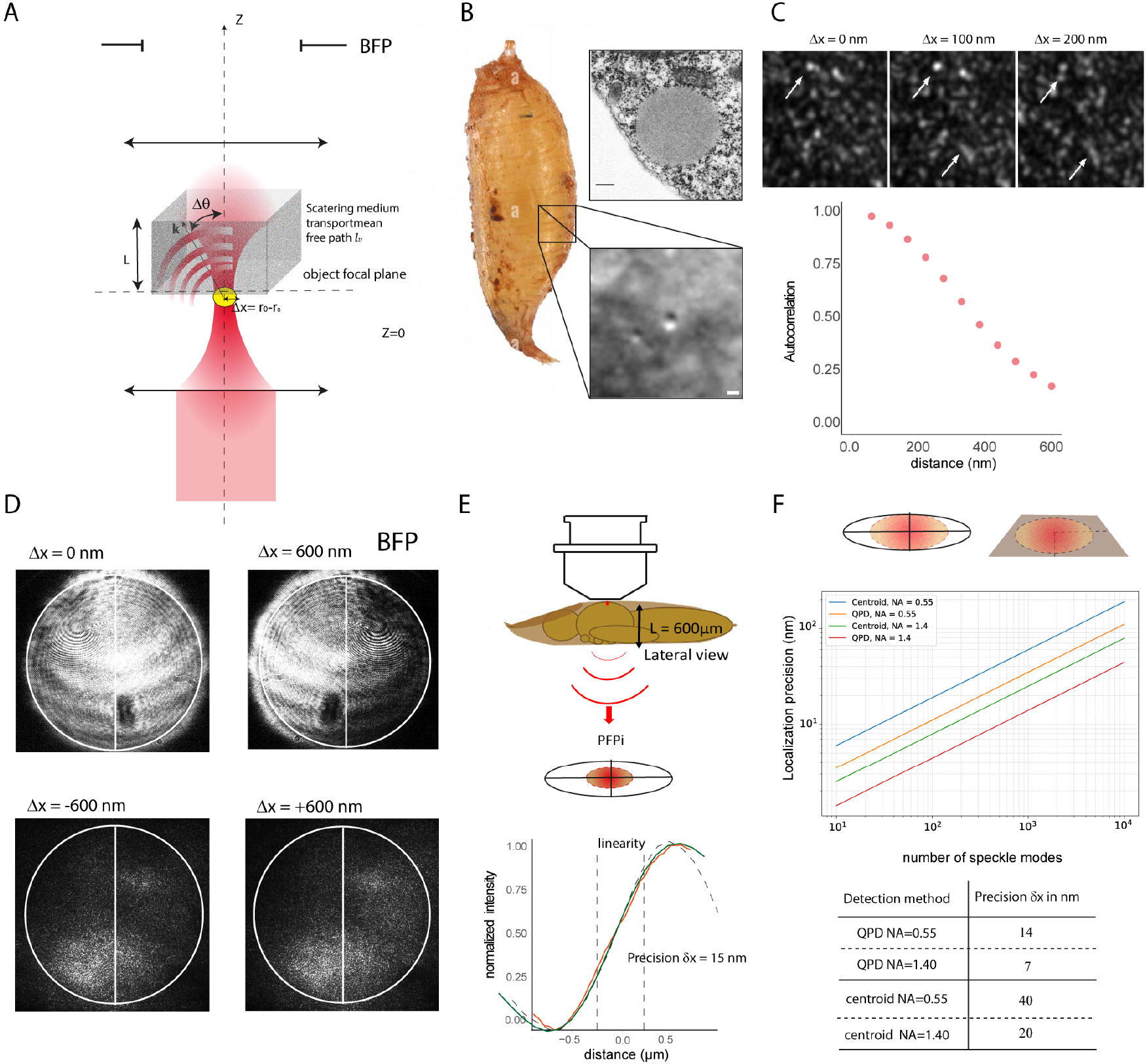
Back focal plane interferometry through turbid biological media enabled by the optical memory effect. (A) A schematic representation of back focal plane interferometry (BFPi) through a thick, optically diffusive biological tissue is presented here. The field detected at the back of the focal plane is the result of the interference between the focused beam that has been incident on the system and the field scattered k by the object which is trapped. These both propagate through the same inhomogeneous medium. The focused beam is close to the surface. (B) Experimental demonstration of the optical memory effect in Drosophila pupa. Lipid droplets are spherical objects that scatter light and produce interference at the back focal plane of the pupa. A typical lipid droplet is shown in an image of TEM (scale bar 200 nm) and a brightfield image (scale bar 1 µm). (C) Top,Intensity images recorded at the image plane of the collection objective for different lateral positions of the focused beam exhibit similar speckle patterns translated according to the beam displacement. Bottom, spatial correlation of the transmitted intensity as a function of beam displacement, defining the memory effect range over which the transmitted field remains correlated. In the pupa, correlations persist for displacements up to approximately 200–300 nm. (D) The intensity image function of the *δX* in the back focal plane, as collected by an optical condenser (NA = 0.55) through water (top) or the living of a Drosophila pupa (bottom). (E) Back focal plane (BFP) interferometric detection in the *Drosophila* pupa using a quadrant photodiode (QPD). The normalized response follows the expected cosine dependence (dashed line), and remains well preserved in the pupa (orange) compared to water (green), despite increased scattering. The signal retains a linear regime suitable for precise displacement measurements, with a localization precision of ∼ 15 nm at a depth of *L* ∼ 600 *µ*m. (F) Localization precision as a function of the number of speckle modes for two numerical apertures (NA = 0.55 and NA = 1.4), comparing centroid-based detection and quadrant photodiode (QPD) interferometry. Increasing speckle contributions degrade the interferometric signal and reduce precision, while higher NA improves sensitivity. QPD detection consistently outperforms centroid estimation due to its differential robustness to speckle noise.

In general, two major approximations are used in the literature to describe this interference pattern. First, it is usually assumed that the trapped droplet is small enough for its radiating pattern to be similar to that of a dipole[21]. Note that more precise calculations using Mie scattering have also been proposed for 3D BFPi [22].

In our study, the incident field is a tightly focused beam with a wavelength of ∼ 1 *µ*m, generated by a high numerical aperture objective (NA = 1.4 or NA = 1.25). In contrast, the collection objective has a lower numerical aperture (NA = 0.55 or up to 1.4, depending on the configuration). The trapped probes—either silica beads or endogenous lipid droplets—are approximately spherical (Fig. 1B, cryoelectron microscopy) and have diameters ranging from 0.3 to 0.8 *µ*m, comparable to or smaller than the illumination wavelength. In this configuration, the trapped objects are located near the surface of a thick, optically scattering biological tissue of thickness *L*. The incident beam is focused at the lower interface of the tissue by the high-NA objective, while the transmitted field propagates through the heterogeneous medium before being collected by the detection objective (Fig. 1A). Importantly, the focal planes of the illumination and collection objectives do not coincide. The illumination objective is focused near the bottom of the scattering layer, whereas the collection objective is effectively conjugated to the top surface of the tissue. This spatial separation defines the interferometric detection geometry in the presence of optical scattering.

We denote by *z* the coordinate along the optical axis, with *z* = 0 corresponding to the focal plane of the illumination objective, and by **r** the transverse coordinate. In our model, both the focus of the incident beam and the center of the scatterer (the droplet) are initially located at *z* = 0, with transverse positions denoted by **r**_0_ and **r**_*s*_, respectively. During tracking, the beam focus is displaced relative to the probe, such that **r**_0_ ≠ **r**_*s*_. For simplicity, we model the angular field distribution of the incident focused beam at the back focal plane (BFP) of the collection objective as that produced by an effective dipole located at (**r**_0_, 0). This approximation remains valid provided that the incident beam is not significantly distorted by aberrations or scattering along its propagation from the laser source to the focal region.

In addition, we model the angular distribution of the field scattered by the trapped probe by that of a dipole placed at (***r***_*s*_, 0). Even though the probe size may reach half the wavelength, this dipole approximation is reasonable within the modest angular range of the collection objective.

Assuming scalar approximation, we introduce the Fourier transform of the Green function *Ĝ* such that *Ĝ*(*k, r*) is the field radiated at a point ***k*** of the Back Focal Plane (BFP) of the collection objective by a dipole placed at a position (***r***, *z* = 0) (***k*** is the wavevector transverse projection of the field plane wave expansion). Importantly, *Ĝ* accounts for the random medium (the biological tissue) and the optical aberrations that are between the dipoles and the BFP (Fig. 1A). With this notation, the total field detected at ***k*** in the backfocal plane satisfies:

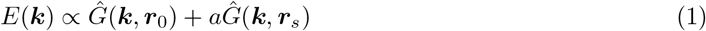

where *a* is a complex number that accounts for the polarizability of the droplet (related to its size and dielectric contrast). For a spherical particle of radius *R* smaller than the wavelength, the scattered field can be described by an effective dipolar response. The polarizability scales as

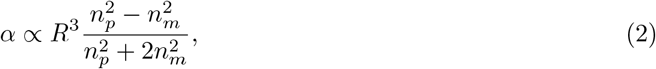

where *n*_*p*_ and *n*_*m*_ are the refractive indices of the particle and the surrounding medium, respectively.

The key point of our model is to assume that the distance between the two dipoles lies within the range of the shift memory effect [23].

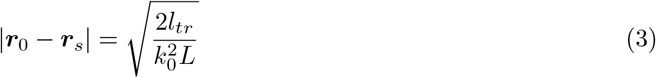

where *L* is the thickness of the heterogeneous layer between the trap and the collection objective, *l*_*tr*_ is the mean free path of the random medium, and *k*_0_ = 2*π/λ* where *λ* is the incident wavelength. Under this assumption, the far-field (equivalently the field measured at the back focal plane of the collection objective) radiated by the two dipoles verifies

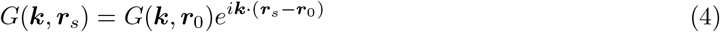

Equation 4 means that, at the image plane of the collection objective (namely at the output of the heterogeneous layer), the field scattered by the droplet is a translated version of the field of the incident focused beam. To test if BFPi is compatible in strong regime of optical scattering, we have measured responses through 600 µm thick *Drosophila* pupae centered on a 800 nm lipid droplet. The sample were moved along the x-axis using a high precision piezoelectric stage (see Supplementary Material). In Fig. 1C we display three images for three different positions of the focused beam. A strong decorrelation is obtained after *δ*_*x*_= 300 nm related to the loss of the memory effect [24]. In Fig. 2C we also plot the correlation between the image obtained when the focus is placed at the origin and the image obtained when the focus departs from the origin, *f* (Δ*x*) = [∫*I*(***x***, 0)*I*(***x***, Δ*x*)d***x*** − ∫ *I*(***x***, 0)d***x*** ∫ *I*(***x***, 0)d***x***]/ ∫ *I*^2^(***x***, 0)d***x*** where *I*(***x***, *a*) is the intensity recorded at the image plane of the collection objective at point ***x*** for a focus position placed at 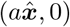.

**Figure 2:**
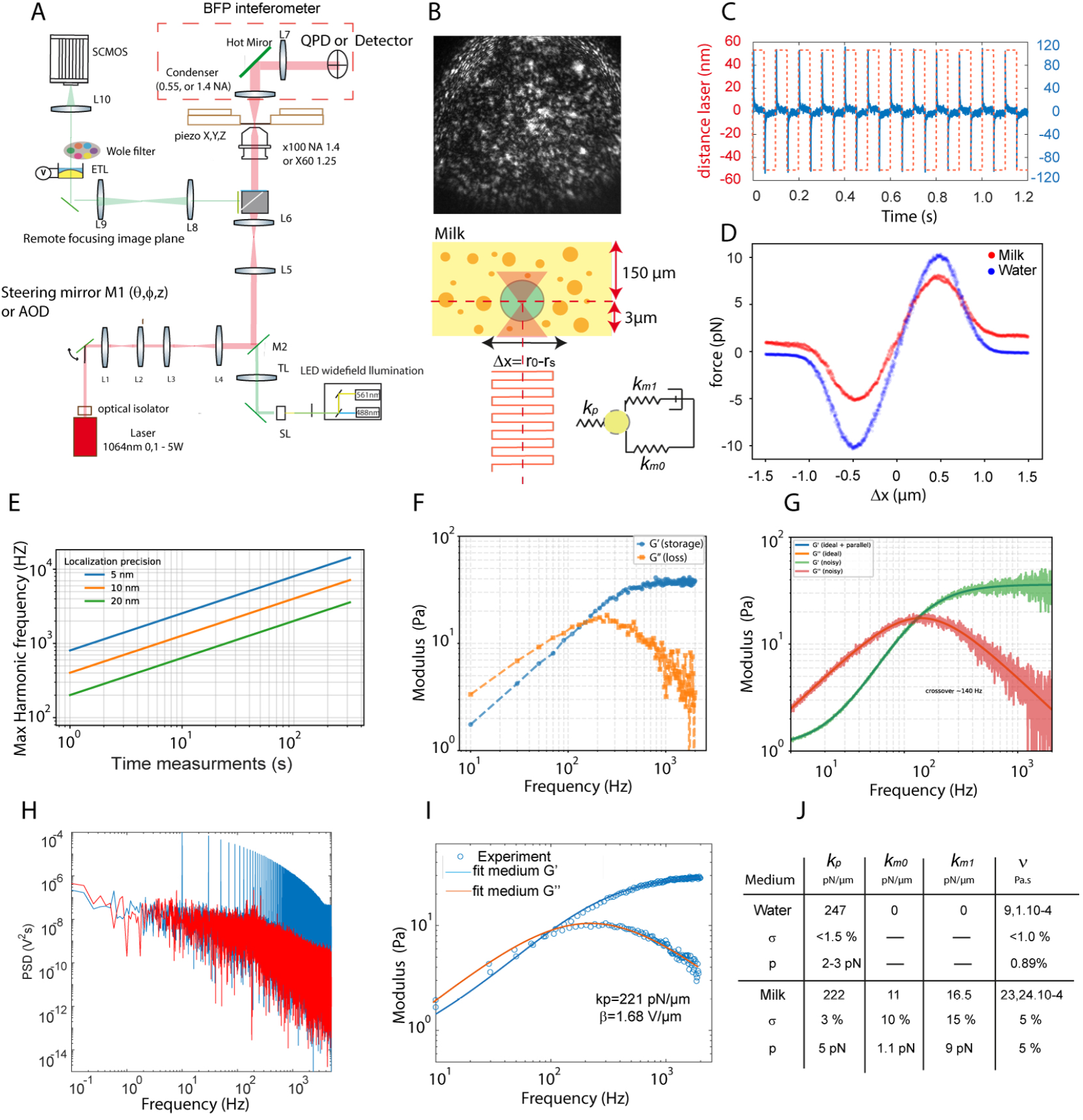
(A).Optical tweezers setup based on back focal plane interferometry (BFP). The detection can be performed either using a quadrant photodiode or centroid tracking from camera images. Beam steering is achieved via either a fast steering mirror or an acousto-optic deflector (AOD), enabling square-wave excitation of the trap position. The interferometric detection is performed in the back focal plane of the condenser objective.(B) Representative imaging field in milk. A trapped bead (1 µm diameter) is positioned within a millimeter-thick viscoelastic medium. The square excitation protocol consists of alternating trap displacements (x 10–15 nm at the bead level), enabling harmonic decomposition of the mechanical response.(C) Example time trace of trap command (red dashed line) and BFP bead response (blue) during square-wave excitation. (D) Rapid spatial force profile of the bead displacement along the trap axis in milk compared to water. (E) Maximum usable harmonic frequency as a function of measurement duration and localization precision (5–20 nm). (F) Measured storage and loss moduli, G and G, of milk obtained from harmonic analysis. The crossover frequency (140 Hz) marks the characteristic relaxation timescale of the medium.(G) Numerical simulation of G and G including phase and localization noise, reproducing the experimental uncertainty envelope.(H) Harmonic power spectral density illustrating the relative contribution of active harmonics and thermal background.(I) Global fit of active and passive spectra using the generic viscoelastic model (Maxwell branch + weak parallel elasticity + fractional contribution), allowing simultaneous extraction of trap stiffness kt, medium elastic parameters,friction coefficient, and detector conversion factor. (J) Extracted mechanical parameters for water and milk.

The correlation function shows that, in the *Drosophila* pupa, the memory effect still holds for displacements up to 200 nm which is generally bigger than the displacement of the trap. In that case, the intensity in the Fourier plane of the optical condenser can be written as follows:

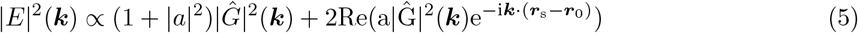

To pursue the analytical calculation, we now assume that *Ĝ* can be considered an uncorrelated zero-mean gaussian random variable. This assumption amounts to considering that the field radiated by a source after transmission by the random heterogeneous sample is a fully developed speckle. It exhibits a homogeneous variance < |*Ĝ* |_2_ >, in which, invoking the ergodicity of the random process, <> corresponds to the spatial averaging of *Ĝ* over enough large domains *s* in the pupil plane.

We now calculate the signal that is obtained by integrating |*E*|^2^ over *I* domains *D*_*i*_, *i* = 1…*I* of the pupil plane (taking for example a four quadrants detector). The key point is to notice that |*Ĝ*|^2^ varies much more rapidly with ***k*** than cos(−*i****k*** · (***r***_*s*_ − ***r***_0_)) because |***r***_*s*_ − ***r***_0_| << *λ*. Thus we can discretize the integration domain into small *s* patches over which the cosine is assumed to be constant, leading to,

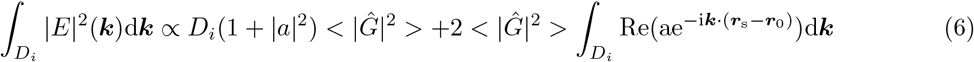

Thus, we retrieve the same expression for the integrated signal as that obtained when the trap is done in a homogeneous medium. In the simple case where the displacement is done along the *x* axis, 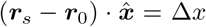 and the pupil is cut in two halves (*D*_+_, *D*_−_) along the *x* axis (see Fig. 1D), we find easily (under small displacement approximation, Δ*x* << *λ*) that,

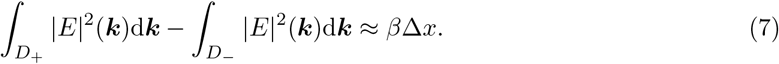

Just prior to applying a continuous displacement setpoint along the *x*-axis, the trapped bead was precisely aligned with the laser focus using the position detectors. This alignment is straightforward to verify, as the *y*-component of the signal must vanish while the *x*-component remains symmetric and centered around zero in the back focal plane (BFP). However, in scattering conditions, the expected cosine dependence of the BFP interferometric signal is distorted by speckle arising from optical heterogeneities at the pupil plane. The degree of speckle decorrelation is related to the photon mean free path *L* within the sample. Autocorrelation analysis shows that displacements on the order of ∼ 300 nm remain within the optical memory effect regime. Figure 1D presents the interference patterns recorded in the pupil plane for identical displacements *δx*, both in the absence and presence of the *Drosophila* pupa. The white regions indicate the effective positions of the detector. Figure 1E compares the experimental BFP response with the theoretical prediction derived in the previous section (Eq. 5). The dashed curve corresponds to the ideal cosine response, while the green and orange curves represent measurements in water and through the *Drosophila* pupa, respectively. Despite increased scattering, the interferometric response measured through the pupa closely follows the expected behaviour, with only moderate noise relative to the water condition. Importantly, a linear response regime is preserved over displacements of a few hundred nanometres (Fig. 1E). This demonstrates that differential BFP measurements remain robust even in strongly scattering environments, enabling reliable tracking of lipid droplets under such conditions.

Building on the previous expression of the interferometric intensity in the presence of scattering (Eq. 5), we now evaluate its impact on localization precision. In this framework, the displacement information remains encoded in the cosine term, but its amplitude is modulated by the speckle-dependent factor |*Ĝ*(**k**) |^2^, which introduces spatial fluctuations across the pupil.

When integrating the signal over the detection aperture, these fluctuations do not fully average out. Instead, the interferometric contrast is effectively reduced by the finite number of statistically independent speckle grains contributing to the signal. Denoting this number as *N*_speckle_, the resulting contrast scales as

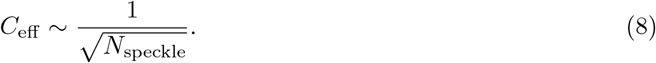

This reduction in contrast directly impacts the slope of the interferometric response with respect to displacement, thereby degrading the achievable localization precision. The displacement uncertainty can be expressed as

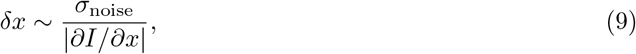

showing that speckle-induced contrast loss leads to an increased uncertainty on position measurements.

The displacement uncertainty *δx* is given by the ratio between the noise amplitude and the slope of the interferometric signal,

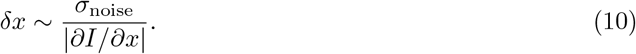

Since both the signal slope and the noise scale with the effective contrast *C*_eff_, the localization precision in a turbid medium obeys

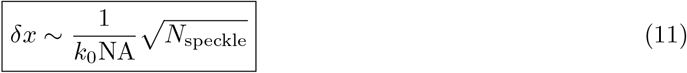

This expression shows that the spatial resolution of BFP interferometry necessarily degrades with increasing speckle complexity, independently of the detection strategy.

In both quadrant photodiode (QPD) and centroid-based detection, the displacement information originates from the same interferometric term from equation (11). The difference between the two detection strategies lies in the way this modulated cosine signal is spatially integrated over the pupil (Fig.1F). The precise details of the calculation of *δx* can be found in the Supplementary Material.

Both detection schemes exhibit the same fundamental 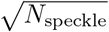 degradation of spatial resolution imposed by speckle-modulated interferometric contrast. However, the QPD performs a differential measurement that suppresses common-mode speckle fluctuations and preserves the global cosine slope, whereas centroid detection amplifies local speckle-induced intensity variations. This explains why, in strongly scattering media, QPD-based BFP interferometry consistently outperforms centroid-based detection, even though neither method can overcome the fundamental contrast loss induced by speckle. The curve end the table in figure 1 -F, shown the precision accuracy depending of the number of speckle grain observe in the backfocal plane. In consequence,For a 1 µm bead with NA=1.4 and *N*_modes_ ∼ 300, the positions detector accuracy have precision *δx*_centroid_ ∼ 20 − 40 nm, roughly an order of magnitude worse than the QPD detector. While the speckle-induced degradation fundamentally limits the spatial resolution of back-focal-plane interferometry in turbid media, the differential nature of quadrant photodiode detection preserves nanometric tracking accuracy. This robustness enables reliable active calibration of both the interferometer and the optical trap stiffness, as developed in the following section using square-wave fluctuation–dissipation measurements.

## 3 Square Wave absolute optical force nanoscopy

### 3.1 methodological principle

When the optical properties of the surrounding medium are unknown or spatially heterogeneous, calibration of optical tweezers cannot rely on simplified hydrodynamic assumptions. This framework builds on the foundations of passive and active microrheology, where the frequency-dependent viscoelastic moduli are inferred from particle fluctuations and response functions, as originally established by Mason andWeitz [25]. In such cases, calibration can be performed using the fluctuation–dissipation theorem (FDT), which relates the thermal fluctuations of a trapped particle to its linear mechanical response [18, 19]. This framework, commonly referred to as FDT optical tweezers, combines passive and active microrheology in a self-consistent manner. In the passive step, the thermal motion of the optically trapped particle is recorded to obtain its power spectral density. In the active step, the trap position (or equivalently the sample stage) is oscillated transversely while the particle displacement is measured with a backfocal plane interferometre expose in part 2. Repeating this procedure across frequencies yields the frequency-dependent response function of the system.

By exploiting the fluctuation–response relationship imposed by FDT, unknown parameters — including trap stiffness *k*_*T*_, viscous drag coefficient Γ, detector conversion factor*β*, and the viscoelastic properties of the surrounding medium *G*^*′*^ *G*^*′′*^ — can be simultaneously determined from the combined passive and active datasets[26]. Laser-driven FDT calibration is particularly robust because trapping and back focal plane interferometric detection are performed simultaneously, enabling high-frequency measurements (up to several hundred hertz) with minimal perturbation amplitude. At sufficiently high frequencies, active nonequilibrium processes such as actomyosin activity become negligible, ensuring that the FDT-based calibration remains valid. To further accelerate spectral acquisition while reducing phototoxic exposure, we extend this framework using a multiplexed square-wave excitation strategy [27] This approach simultaneously probes multiple harmonics within a single acquisition window, increasing spectral efficiency and reducing total measurement time. The full formalism, including the differential BFP interferometric signal, active and passive spectra in a viscoelastic medium, and parameter extraction equations, is provided in the Supplementary Information.

A key advantage of square-wave driving is that the imposed trap displacement is known with nanometric accuracy, independently of the optical properties of the sample.As a result, the conversion factor *β* between the BFPi signal and physical displacement can be calibrated directly for each individual measurement, even in strongly scattering environments. This point is critical in living tissues, where optical diffusion alters the absolute intensity distribution in the pupil plane but preserves the linear interferometric response Low-frequency square steps are used to probe quasi-static mechanical responses, while high-frequency square-wave modulation (up to several hundred hertz) is employed for FDT-based calibration, where active cellular processes are negligible.

Then, thanks to the Onsager principle [28], the optical trap for each frequency can be written as the ratio between the passive and active spectrum for each harmonics :

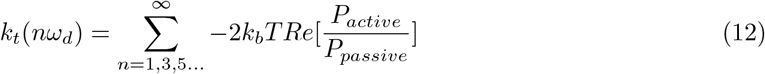

where *k*_*t*_ the stiffness of the optical tweezers, *k*_*b*_ the Boltzmann constant, *T* is the temperature, *P*_*active*_ and *P*_*passive*_ are the active and passive powers, respectively.

In complex viscoelastic media, the bead does not experience the optical trap stiffness *k*_*T*_ alone, but rather an effective frequency-dependent stiffness given by

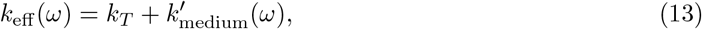

where 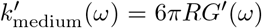 represents the elastic contribution of the surrounding material. In the high-frequency regime of our measurements, *G*′(*ω*) generally increase to 20 200 Pa for a bead of radius *R* = 0.5 *µ*m, this corresponds to an additional stiffness of ∼ 165 pN*/µ*m. For a trap stiffness of ∼ 250 pN*/µ*m, the viscoelastic contribution therefore represents more than 60% of the trap stiffness. If one performs a FDT conventional calibration while neglecting the medium elasticity, the extracted stiffness corresponds to *k*_eff_ rather than *k*_*T*_. This leads to a systematic overestimation of the true optical trap stiffness. The magnitude of this bias depends on the frequency window used for calibration and becomes particularly significant when measurements extend beyond the viscoelastic crossover frequency. For this reason, we adopt a two-step strategy. First, we perform an independent, conventional calibration to obtain an initial estimate of the trap stiffness and detector sensitivity *β*. Second, we refine these parameters through a global fitting procedure in which the trap stiffness *k*_*T*_, interferometric conversion factor *β*, friction coefficient *γ*, and viscoelastic parameters of the medium are fitted simultaneously with a generic viscoelastic model explained in detailed in supplementary method. This global approach consistently accounts for the mechanical coupling between the trap and the viscoelastic environment.

Importantly, the global method avoids parameter bias arising from partial modeling of the system and ensures that the extracted trap stiffness corresponds to the intrinsic optical confinement rather than to the combined trap–medium response. This is particularly crucial in soft viscoelastic systems where the medium elasticity is comparable to the trap stiffness.

The accessible frequency window and parameter precision in square-wave interferometric microrheology are fundamentally governed by localization accuracy and acquisition time. A full derivation of the uncertainty scaling, harmonic signal-to-noise limits, and phase error propagation is provided in the Supplementary Material. In summary, a square-wave excitation of amplitude *A* and localization precision *σ*_*x*_, the signal-to-noise ratio of the *n*-th harmonic scales as

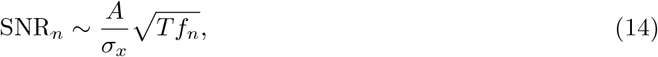

where *T* is the measurement duration and *f*_*n*_ the harmonic frequency. Consequently, the maximum usable frequency increases with acquisition time and improves quadratically with localization precision define in section 2. In practice, nanometric tracking enables extension of harmonic analysis into the kilohertz regime within seconds of acquisition.

The relative uncertainty on the trap stiffness and on the extracted viscoelastic moduli follows the same statistical scaling:

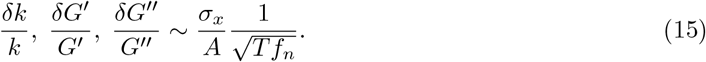

Thus, precision is primarily limited by displacement accuracy, but can be improved through statistical averaging over harmonic cycles. Higher harmonics naturally benefit from increased sampling density, which provides enhanced robustness, even in the presence of low-frequency active fluctuations.

Finally, the phase shift between the applied force and the bead displacement directly reflects the balance between elasticity and dissipation :

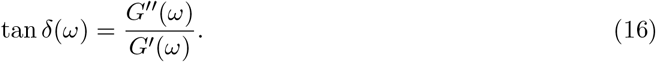

Since harmonic phase is extracted independently at each frequency, the global fitting procedure intrinsically accounts for elastic contributions from the surrounding medium. Crucially, phase uncertainty is bounded by localisation precision and does not fundamentally restrict spectral reconstruction once a sufficient harmonic signal-to-noise ratio has been achieved.

### 3.2 Experimental implementation end validation of squarewave FDT optical tweezer in complex viscoelastic turbid media

Both optical tweezers systems used in this study combine a fluorescence microscope with an active optical trapping module. They rely on back focal plane (BFP) interferometric detection, which provides sub-nanometre displacement sensitivity and enables broadband force measurements. Two complementary implementations were employed: a custom-built setup and a commercial system from Impetux.

These platforms differ in several key experimental aspects, including the detection strategy (quadrant photodiode versus position sensor), the laser beam steering approach (kHz-bandwidth steering mirror versus high-frequency acousto-optic deflector), and the overall acquisition bandwidth (25 kHz versus 1 kHz).A simplified representation of the two optical configurations is shown in Fig. 2a, while full technical details of both setups are provided in the Methods section. A 1064 nm laser beam is focused through a high numerical aperture objective to generate the optical trap (NA 1.4 ×100 oil objective in the homemade system and NA 1.25 ×60 water immersion objective in the Impetux system). In the homemade setup, the forward-scattered and unscattered light are collected by a condenser objective and projected onto the detection plane. A condenser with moderate numerical aperture (NA = 0.55) is used, which facilitates sample accessibility while preserving interferometric detection and three-dimensional tracking capability. The commercial Impetux system relies on optimized high numerical aperture optics (NA = 1.4) designed for precise force measurements and position detection. Depending on the experimental constraints—such as bandwidth requirements or imaging depth—two detection strategies can be employed: wideband quadrant photodiode detection, providing kHz-range interferometric measurements, or centroid-based position tracking at up to 25 kHz, which is advantageous in strongly scattering samples where interferometric contrast may be reduced.

The trapping laser power delivered to the sample ranged from 50 to 200 mW. At this wavelength and power level, no detectable alteration of tissue behaviour was observed, with an estimated temperature increase limited to 0.3–1.5 °C. In our experiments, beads or lipid droplets were trapped at depths ranging from 3 to 20 *µ*m above the coverslip, after which the square-wave FDT calibration protocol was applied.

The method was first validated in a controlled scattering medium using milk and calibrated 1 *µ*m polyester beads (*n* = 1.56). Milk is a non-Newtonian fluid composed of casein micelles (50–300 nm), which induce strong optical scattering. Measurements were performed at a depth of ∼ 150 *µ*m, where speckle formation is significant (Fig. 2B). Such media are well described by casein-based emulsions exhibiting Maxwell-like behaviour with distributed relaxation spectra [29, 30, 31, 32, 33], providing a relevant testbed for tissue-like mechanical conditions. We first used the commercial Impetux system to probe the high-frequency square-wave FDT response in milk and to compare it with direct momentum-based force measurements [17]. The bead response shows both amplitude attenuation and phase lag relative to the imposed 100 nm square-wave excitation (Fig. 2C), reflecting the complex viscoelastic response of the medium. In contrast to sinusoidal excitation, the square-wave approach enables simultaneous probing of multiple harmonics within a single acquisition window, thereby improving spectral efficiency. The fast response of the acousto-optic deflector (AOD) further allows rapid estimation of the interferometric response and trap stiffness. A direct comparison of bead interferometric response profiles in water and milk (Fig. 2D) at 10 mW reveals a marked reduction in the interferometric amplitude in milk. This difference cannot be explained solely by the small refractive index mismatch between water (*n* = 1.33) and milk (*n* = 1.348), indicating a dominant contribution of optical scattering to the measured response. Consistently, the direct momentum method yields *k*_*p*_ = 28.9 pN*/µ*m in water and *k*_*p*_ = 19 pN*/µ*m in milk at 10 mW, highlighting the strong impact of scattering on force measurements performed with high numerical aperture optics.

The FDT square wave optical method is then applied at 100 mW in the same bead to study the validity of our model and the sensitivity in the 2.5 kHz frequency range. The harmonic power spectrum (Fig. 2H) confirms that active harmonics dominate over the thermal background within the usable frequency window. Because the viscoelastic contribution can approach the trap stiffness, independent trap calibration may lead to systematic bias if the medium elasticity is neglected. As outlined in the Supplementary Material, the maximum usable harmonic frequency is contingent on two factors: the precision of localisation and the total acquisition time (Fig. 2E). This protocol generates a discrete harmonic spectrum at odd multiples of the fundamental excitation frequency. For a localization accuracy induced by the Milk of 5–20 nm and acquisition times ranging from seconds to minutes, harmonic analysis can extend into the kilohertz regime. We therefore implement a global fitting approach, combining active harmonic response and passive fluctuation spectra within a unified viscoelastic model. Thanks to the exact knowledge of the beads size from the fast scanning with the AOD end the estimation of the optical trap stifness (kp) from the global approach (Fig. 2I), the measured storage and loss moduli, *G*′(*ω*) and *G*″(*ω*), exhibit a characteristic crossover frequency near 2Khz (Fig. 2F), consistent with a dominant relaxation timescale on the order of milliseconds. The high-frequency plateau of *G*′ (35 Pa) corresponds, for a 1µm diameter bead. This value represents a substantial fraction of the optical trap stiffness (221 pN/µm), indicating that the bead experiences a combined confinement arising from both trap and medium elasticity. To assess measurement robustness, we simulated the propagation of localization and phase noise onto the extracted moduli (Fig. 2G) from equation explained in Supplementary Material. The uncertainty envelope reproduces the experimentally observed increase in relative noise at high frequency, particularly for *G*″, which is more sensitive to phase fluctuations. The agreement between harmonic analysis, passive spectra, numerical simulations, and global fits demonstrates the internal consistency of the square-wave FDT implementation. The method enables absolute sub-piconewton force spectroscopy in thick viscoelastic environments without requiring prior independent calibration in a reference Newtonian fluid. This global strategy ensures consistent extraction of mechanical and instrumental parameters while accounting for the mechanical coupling between trap and medium. The resulting fitted parameters for water and milk are summarized in Fig. 2J. The difference in value between the water and the milk is more realistic compared to the estimation made using direct optical momentum measurements in turbid media, providing a robust framework for quantitative mechanical measurements in complex living tissues. The method can extract in only 10 seconds of measurments : the intrinsic trap stiffness *k*_*T*_, conversion factor *β*,the full spectrum friction coefficient *γ*, the full spectrum of *G*′(*ω*) and *G*″(*ω*) using a Maxwell branch describing the dominant relaxation mode, a weak parallel elastic contribution accounting for residual low-frequency stiffness, and a fractional power-law component capturing distributed relaxation dynamics.

## 4 Revealing subcellular mechanical properties in living tissues

### 4.1 Singe cell microrheology from square-wave har-monics in drosophila leg

We next tested the robustness of the global square-wave FDT calibration in a mechanically and optically challenging environment: endogenous lipid droplets embedded within the curved leg tissue of Drosophila. Brightfield imaging enabled the selection of individual droplets (Fig. 3A), whose diameters were independently validated by electron microscopy and back focal plane interferometry (Fig. 3C), ensuring accurate probe size determination. A fast scan using AOD excitation produced well-defined interferometric harmonic responses across droplets of varying size (Fig. 3B). For this tissue, the optical scattering is lower, with only hundreds of speckle modes observed in the Fourier plane of the condenser. The scanning distance can then be 1.5 µm without loss of correlation of the speckles. However, trap stiffness estimates obtained via the conventional momentum-based calibration exhibited strong sensitivity to optical aberrations induced by tissue curvature, leading in some cases to overestimations of *k* exceeding 50 (Fig. 3D). These aberrations alter the detected momentum flux without reflecting a true change in optical restoring force, thereby biasing the calibration. In contrast, the square wave (1 HZ with 100nm amplitude) global approach simultaneously fitting passive fluctuations and active harmonic responses remained stable under identical optical conditions. The experimentally measured real and imaginary parts of the response function closely matched the model reconstructed from the global fit (Fig. 3E–F), indicating accurate capture of both elastic and dissipative components of the bead dynamics.

**Figure 3:**
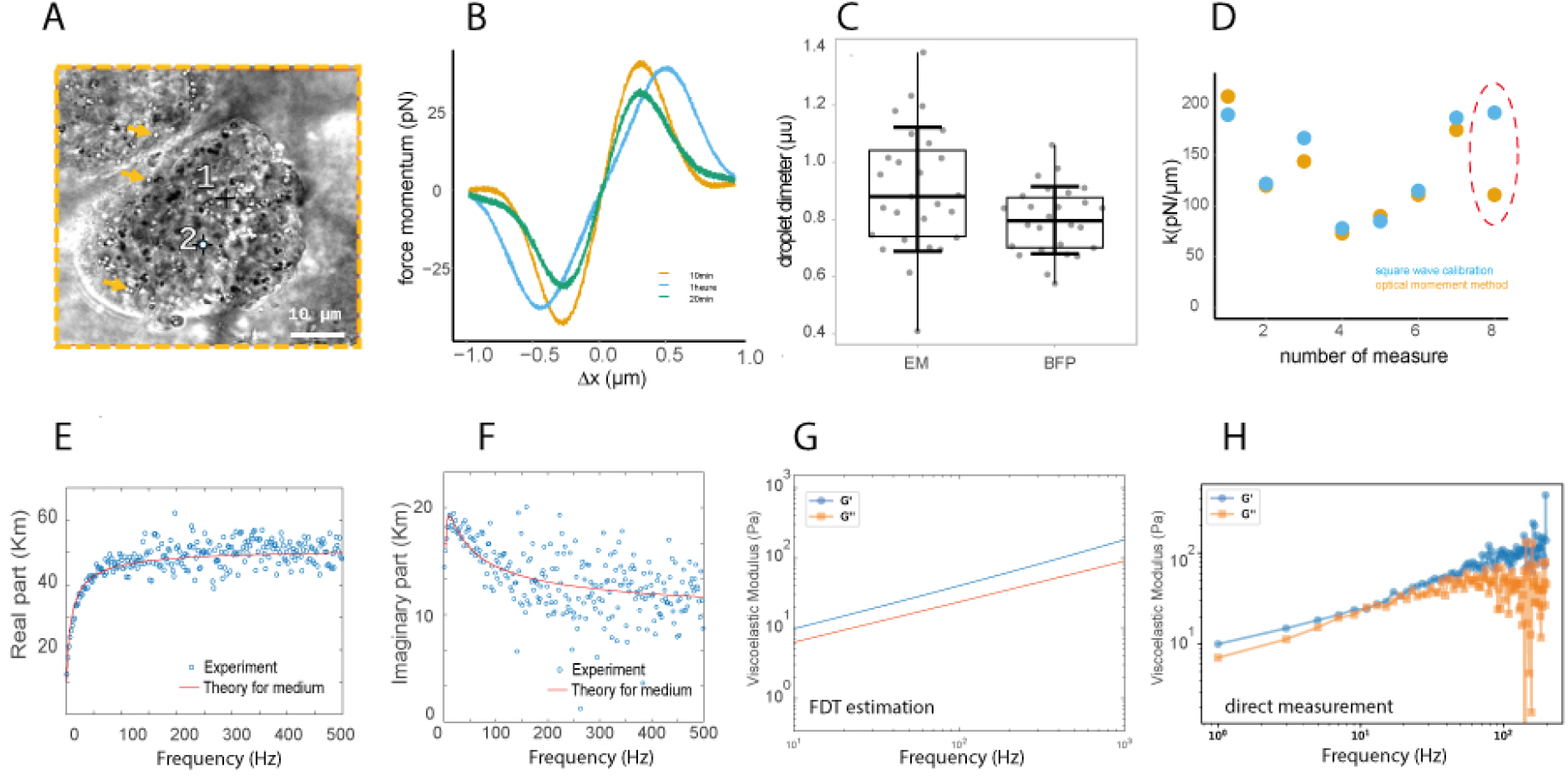
Global calibration enables robust viscoelastic measurements in curved Drosophila leg tissue.(A) Brightfield image of a Drosophila leg showing endogenous lipid droplets used as probe particles. Two representative droplets selected for interferometric microrheology are indicated. Scale bar, 10 µm. (B) Examples of interferometric force–displacement responses under square-wave excitation for droplets of different diameters, illustrating reproducible harmonic behavior within the tissue. (C) Droplet diameters independently measured by electron microscopy (EM) and back focal plane interferometry (BFP), demonstrating consistent size estimation across modalities. (D) Trap stiffness *k* obtained from repeated measurements using the global square-wave FDT calibration (blue) compared with the conventional momentum-based method (orange). In the presence of optical aberrations induced by tissue curvature (red dashed region), the moment method yields substantial overestimation of *k*.(E–F) Real (E) and imaginary (F) parts of the complex response function measured experimentally (points) and reconstructed from the global fit (solid lines), showing excellent agreement across frequencies.(G) Storage and loss moduli *G*′(*ω*) and *G*″(*ω*) extracted directly from the global fit parameters. (H) Independent modulus determination using the corrected trap stiffness obtained from the global calibration. The close agreement with panel (G) validates the consistency and robustness of the global approach. In contrast, using the stiffness estimated by the moment method (not shown) would lead to systematic errors approaching 50

The viscoelastic moduli *G*′(*ω*) and *G*″(*ω*) extracted from the global fit parameters (Fig. 3g) were further validated by an independent modulus reconstruction using the corrected trap stiffness obtained from the same global calibration (Fig. 3h). The close quantitative agreement between the two determinations confirms the internal consistency of the method. Importantly, had the stiffness derived from the moment-based approach been used, the resulting moduli would have been systematically biased by up to 50pc, highlighting the necessity of global calibration in heterogeneous and optically distorted biological tissues.

Having established the robustness of the global calibration framework in the optically challenging and mechanically heterogeneous environment of the Drosophila leg, we next considered its applicability under biologically active conditions. The leg measurements demonstrate that reliable stiffness and modulus extraction can be achieved without relying on high-bandwidth detection or large numerical aperture collection, even in the presence of curvature-induced aberrations. Importantly, the mechanically informative spectral window was found to remain confined below 300 Hz, beyond which phase conditioning deteriorates and the response becomes either elasticity- or drag-dominated. This observation justifies the use of a simplified detection architecture for subsequent in vivo experiments, consisting of a 1 kHz acquisition system and a quadrant photodiode with moderate numerical aperture (NA = 0.55). At this sampling rate (Nyquist frequency 500 Hz), the entire mechanically relevant frequency range is fully captured, while avoiding unnecessary high-frequency noise amplification. Extending the bandwidth beyond 300 Hz would not improve mechanical inference, as the information content is fundamentally limited by mechanical and thermal constraints rather than detector speed.

### 4.2 Cortical tension properties of epithelial cells in drosophila pupa during development using simplify home made setup

We therefore applied this configuration to cortical tension measurements in 600µm thick Drosophila pupae, including strong optical scattering and optical abberration link to the geometry of the sample. We performed absolute force measurements in the epithelial tissue of Drosophila pupa at different developmental stages using the homemade setup Fig.4A. All measurements were performed in intact Drosophila pupae at 3-5µm from the cover slide with high numerical microscope lense (X100 NA=1.4), end interference formation is perturbed by the same strongly turbid optical conditions exposed in section 1. Based on independent characterization of the back focal plane signal, we estimate that approximately *N* ≃ 10^3^ speckle modes contribute to the detected interferometric signal at the pupil plane. Under these conditions, the back focal plane interferometry signal remains detectable but exhibits a reduced interference contrast due to speckle-induced phase scrambling. Optical trapping was achieved using a 1064 nm laser with an injected power of 90 mW at the back pupil of a high numerical aperture objective (NA = 0.55) from the home made setup with a 1Khz band width. Endogenous lipid droplets (typical diameter ∼ 1 *µ*m) were used as force probes. Trap calibration was performed locally for each measurement using the square-wave multiplexed fluctuation–dissipation (FDT) protocol described above. From the active–passive spectral fitting over multiple harmonics, the optical trap stiffness was determined with a relative uncertainty below 10%. Taking into account the reduced interferometric contrast induced by speckle, the effective spatial localization precision of the trapped droplet was estimated to *δx* ≃ 10 nm consistent with independent measurements performed under similar scattering conditions. After calibration of both the detector conversion factor *β* and the trap stiffness *k*_*t*_, slow square-wave displacements of the optical trap were applied in order to probe cortical membrane tension. The trap position was displaced by steps of amplitude Δ*x* = 300 nm at a low frequency of 0.12 Hz, ensuring quasi-static conditions with respect to the viscoelastic relaxation time of the tissue (Fig 4.B). Despite the strong optical diffusion, interferometric tracking remained stable and allowed precise measurement of bead relaxation following square-wave indentation. Representative relaxation curves (Fig. 4,B) exhibit a monotonic decay characteristic of membrane-dominated mechanics. At this low driving frequency, inertial and viscous contributions are negligible, and the mechanical response of the system is dominated by membrane tension. The resulting bead displacement exhibits a rapid elastic response followed by a slow relaxation toward a plateau. In this regime, the force balance reduces to

**Figure 4:**
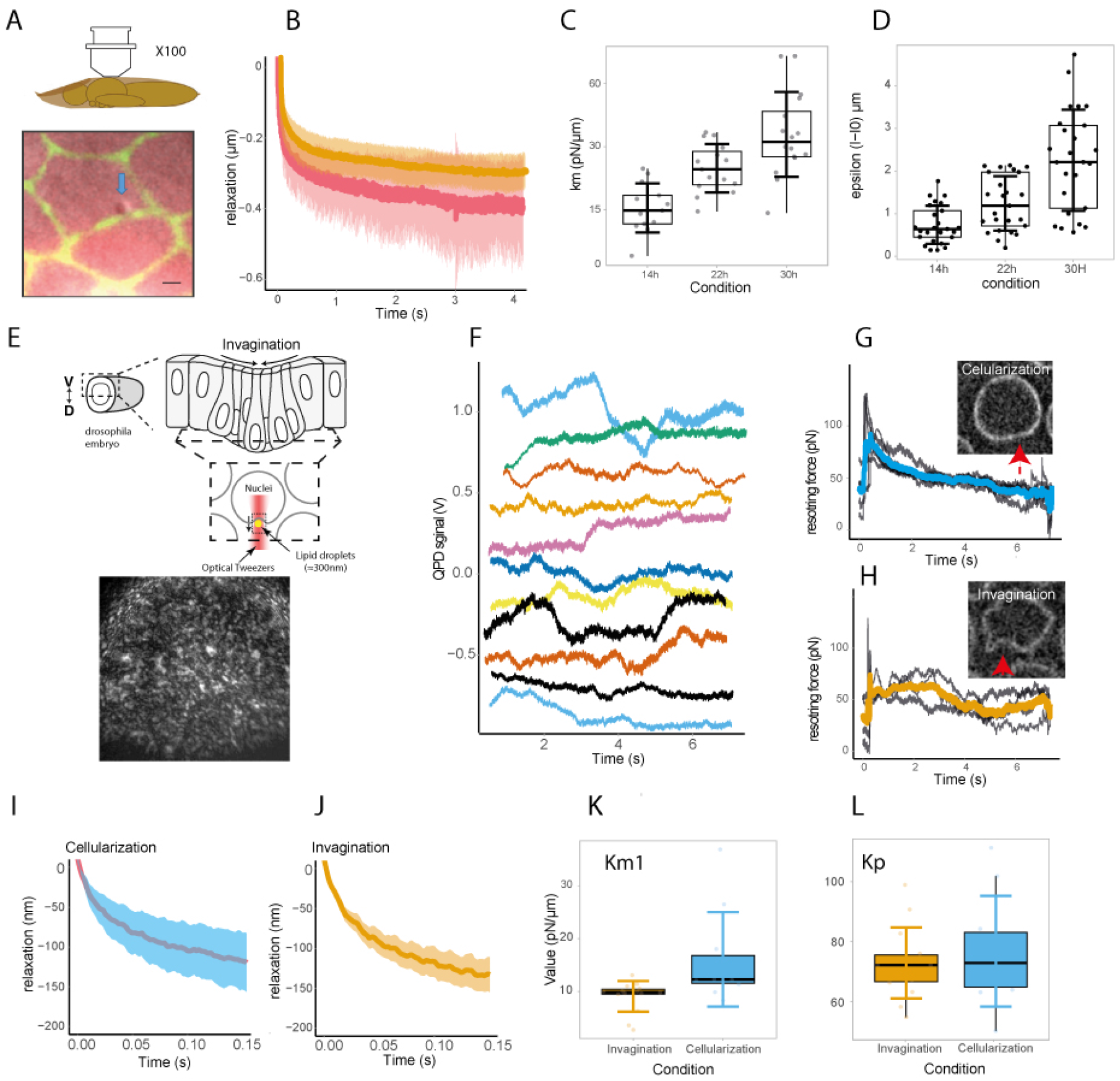
Mechanical measurements in Drosophila pupae and embryos.(A) Overview of the pupal model used to probe cortical mechanics. Lipid droplets are used as mechanical probes to sense the cortical tension of surrounding epithelial cells. Inset: magnified view of a droplet embedded within the tissue (scale bar 1µm).(B) Mean relaxation responses of droplets following mechanical perturbation at two developmental stages (14 h and 30 h after puparium formation). Shaded regions represent the variability across measurements.(C) Distribution of cortical tensions estimated from FDT optical tweezers analysis at different developmental stages (14 h, 22 h, and 30 h). Box plots show the median and interquartile range, with individual points corresponding to single measurements.(D) Independent validation of cortical tension measurements using laser photoablation experiments, confirming the increase in tension during development.(E) Schematic of the experimental configuration used for mechanical measurements in the embryo, together with a representative speckle image used for inter-ferometric detection.(F) Example time traces of quadrant photodiode (QPD) signals recorded during measurements of nuclear fluctuations in the embryo. Each trace corresponds to an individual measurement.(G) Representative images of embryonic nuclei probed in two developmental conditions: during cellularization or during invagination. Red arrows indicate the position of the probe.(H) Examples of noisy indentation responses measured in both conditions (gray traces), together with the corresponding averaged responses (colored curves).(I–J) Mean mechanical responses to high-frequency excitation (2.6 kHz) measured during cellularization (I) and during invagination (J). Shaded regions indicate variability across measurements.(K) Estimated effective nuclear tension Km obtained from harmonic filtering of the response signal.(L) Estimated optical trap stiffness Kp for the two experimental conditions.

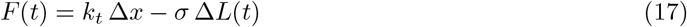

where *σ* is the effective cortical membrane tension and Δ*L*(*t*) the deformation along the pulling direction. The relaxation curves were fitted using a minimal model assuming a single effective membrane tension, corresponding to a plateau force at long times. No additional viscoelastic parameters were required to describe the data within experimental uncertainty. Cortical tension measurements were performed at three distinct developmental stages of the pupa (14H, 22H, and 34H). For each stage, measurmements were repeated on multiple cells, and force–displacement curves were averaged to reduce stochastic intracellular fluctuations. We observe a systematic increase in cortical tension as development proceeds, in quantitative agreement with independent measurements obtained using laser ablation of cell–cell junctions (Fig 4C). In particular, the absolute values of membrane tension extracted from optical tweezers measurements fall within the range inferred from recoil velocities following laser ablation (fig 4D), while providing direct local force calibration. Given the localization precision *δx* ≃ 10 nm and the applied square-wave amplitude Δ*x* = 300 nm, the relative uncertainty on the applied force and extracted membrane tension for each measurments is estimated as

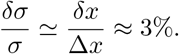

This level of precision demonstrates that, despite strong optical scattering and active intracellular noise, absolute single-cell force measurements with sub-10% accuracy are achievable in intact living tissues using our calibration strategy. The dominant limitation arises from speckle-induced degradation of interferometric contrast rather than from optical trap stability or laser power.

### 4.3 Nuclear membrane tension under active intracellular processes during

We next investigated nuclear membrane tension during mesoderm invagination in the *Drosophila* embryo,a mechanically active system where cells from the ventral part of the embryo constrict their apex thanks to the formation and contractility of a actomyosin meshwork formed below the apical surface of the cell. This constitutes a noisy environment, with strong ATP-driven fluctuations, particularly challenging to measure subtle difference in nuclear membrane stiffness. The optical scattering conditions in the embryo are moderate and correspond to a limited number of speckle modes in the back focal plane, estimated to be on the order of *N*_speckle_ ∼ 10^2^ for a characteristic speckle grain size of ∼ 300 nm (Fig. 4E). Under these conditions, the interferometric contrast of the back focal plane (BFP) signal remains sufficiently high, and the precision of optical tracking is not significantly affected by speckle-induced decorrelation. Because invagination alters the depth of the measurement, it introduces varying optical conditions across experiments, although calibration of the optical trap and position detection was maintained for each measurement.

Measurements were performed under conditions similar to those used for the pupal experiments, with an optical trapping power of 90 mW. Following calibration of the optical trap in the cytosol, controlled indentations were performed on the nuclear membrane using a trapped lipid droplet (Fig. 4G,H). These measurements provide a direct estimate of the apparent mechanical response of the nucleus in two distinct developmental states. During cellularization, nuclei exhibit a stronger restoring force upon indentation, consistent with their more spherical morphology observed in imaging. In contrast, during invagination, nuclei display a reduced restoring response and a visibly deformed contour, strongly suggestive of lower effective tension and/or increased mechanical compliance.

The difference between the two mechanical states is important for validating the subsequent global fitting analysis. Despite strong ATP-driven fluctuations, particularly during invagination, indentation measurements can still distinguish meaningful mechanical differences between conditions. However, active noise significantly reduces their precision at low tension, which diminishes the reliability of low-frequency estimates. These results underscore the inherent limitations of direct indentation approaches in highly dynamic systems, emphasizing the need for harmonic-based global analysis.

Building on this observation, small-amplitude, square-wave displacements with an amplitude of 290 nm and a frequency of 2.6 Hz are applied directly at the nuclear membrane during 17 s. For each condition, the analysis combines 15 independent measurements, each averaged over multiple square-wave periods. This significantly improves the signal-to-noise ratio (see Fig. 4I, J). The resulting relaxation curves are then used as input for the global fluctuation-dissipation fitting procedure, which uses the harmonic content of the response to separate intrinsic viscoelastic properties from active fluctuations.

The global fitting yields robust estimates of both the effective membrane stiffness parameter (*k*_*m*1_) and the trap stiffness (*k*_*p*_) (Fig. 4K,L). Importantly, the extracted values confirm the trend observed in indentation experiments, while providing a significantly improved precision. The consistency of *k*_*p*_ across conditions further demonstrates the stability of the optical calibration, even in heterogeneous and dynamically active environments.

Together, these results demonstrate that, although low-frequency indentation measurements are impacted by ATP-driven activity, harmonic-based global analysis effectively suppresses active noise contributions at higher frequencies. This increases the precision of membrane tension measurements to 4% (see Supplementary Material), enabling reliable and quantitative mechanical characterization of Drosophila embryos.

## 5 Discussion

In this study, we present a square-wave interferometric optical tweezers platform that enables quantitative force spectroscopy in strongly scattering and mechanically active environments, such as developing biological tissues. This approach combines interferometric detection in turbid media with square-wave harmonic decomposition and global fitting of the viscoelastic response. Leveraging the harmonic content of the signal effectively filters active biological noise, enabling robust extraction of intrinsic viscoelastic parameters in regimes where conventional microrheology approaches typically fail.

The angular memory effect and how it constrains interferometric detection is a critical yet often overlooked aspect of deep-tissue optical micromechanics. Precise back-focal-plane interferometry is possible even at millimetric depths thanks to the preservation of phase correlations over small angular ranges in strongly scattering media. However, operating within this regime is not trivial: it requires working within the angular memory window while ensuring there are sufficient ballistic and quasi-ballistic components to sustain coherent interference. Our implementation carefully balances numerical aperture, trapping geometry and detection bandwidth in order to meet these conditions. The robustness observed in the pupa and leg cortex therefore arises from operating in a controlled interferometric regime where, despite multiple scattering, phase information is preserved. Our approach is distinguished from imaging-based methods by this explicit consideration of the optical memory effect, as the latter typically degrade rapidly with depth due to loss of contrast rather than phase coherence.

Unlike classical sinusoidal forcing, the square-wave protocol disperses mechanical energy across harmonics. This enables multi-frequency probing to be performed simultaneously within a single time window. This dramatically improves spectral coverage and robustness in noisy biological systems. Mechanical measurements in developing tissues have been extensively pioneered by groups such as that of Pierre-François Lenne [34], [35], [36],[37],who have demonstrated the importance of cortical tension and cell–cell junction mechanics in morphogenesis using laser ablation, magnetic droplets, optical tweezers or micro-indentation approaches. These methods have provided crucial insights into tissue-scale force balance and tension anisotropy. However, they usually involve geometric relaxation analysis at a low frequency, imaging-based deformation tracking or inferring bulk mechanics without direct force calibration at the nanometre scale. In contrast, the present approach differs in three fundamental ways. First, detection scheme measures bead displacement interferometrically with sub-nanometric precision through turbid media, independent of imaging contrast. This enables measurements in highly scattering samples such as the Drosophila pupa or leg cortex. Optical trap stiffness and force are directly calibrated, allowing extraction of absolute mechanical moduli rather than relative tension proxies. Rather than inferring a single effective tension or stiffness parameter, we extract the full frequency-dependent viscoelastic response. This provides access to dissipation, active contributions, and rheological transitions. Thus, while previous tissue-scale studies have elucidated force balance and morphogenetic tension patterns, our method bridges the gap between cellular biophysics and tissue mechanics by providing local, quantitative, frequency-resolved measurements in intact living tissues.

A central difficulty in living systems is the presence of active, ATP-driven fluctuations that violate fluctuation-dissipation equilibrium assumptions. Our results demonstrate that: low-frequency square-wave relaxation remains suitable for cortical tension estimation, high-frequency harmonic decomposition isolates intrinsic viscoelasticity, harmonics above the fifth order are largely insensitive to collective neighboring cell forces. This harmonic filtering effect constitutes a key conceptual advance. Rather than attempting to suppress active noise experimentally, we exploit the spectral structure of the forcing signal to separate passive mechanics from active fluctuations. This allows quantitative microrheology even in strongly non-equilibrium systems such as the early Drosophila embryo.

Recent work on time-shared optical tweezers for microrheology has demonstrated that, with high throughput, multiplexed traps can measure the age-dependent viscoelastic properties of organelles, cells and organisms. This method offers significant advances in parallelisation, time-resolved ageing studies and long-duration mechanical tracking. These recent time-shared optical microrheology approaches, including TiSOM [38], leverage interferometric detection but rely on transmission-based optical models to infer forces and moments. While these models are powerful for weakly scattering or quasi-homogeneous samples, they implicitly assume stable forward-propagating fields and negligible multiple scattering. In thick or heterogeneous tissues, however, this assumption progressively breaks down, resulting in systematic bias in force reconstruction and mechanical parameter estimation. In contrast, our framework does not depend on a transmission model of the sample. Force calibration remains local and trap-based, and mechanical parameters are extracted through harmonic decomposition rather than direct moment inference. This distinction is particularly important in millimetric, anisotropic or dynamically remodelling tissues, where optical propagation cannot be approximated by a simple transmission kernel. This approach ensures quantitative accuracy in situations where traditional microrheology methods become challenging to apply. Nevertheless, it is chiefly dependent on conventional sinusoidal or stochastic microrheology, frequently under conditions more proximate to equilibrium.

Our strategy is supplementary and differs in a number of ways. Firstly, the typical process of time-shared microrheology involves the reconstruction of mechanical spectra from stochastic fluctuations or sinusoidal driving.In contrast, the square-wave protocol utilises a different approach by encoding multiple discrete harmonics simultaneously. This enables global fitting with intrinsic phase relationships between harmonics.Secondly, in strongly active tissues, stochastic microrheology may be contaminated by ATP-driven fluctuations. So, our harmonic filtering provides a built-in separation between active low-frequency modes and intrinsic viscoelastic response, and our interferometric strategy detection enables operation in highly scattering millimetric tissues.

Integrating time-sharing with our square-wave interferometric approach represents a promising next step. A dual-trap configuration would enable simultaneous active and passive microrheology, spatial correlation measurements, and mapping of mechanical heterogeneity across tissues, while allowing parallel probing of multiple cells during morphogenesis. Such time-shared square-wave protocols could further provide higher-throughput access to the dynamics of viscoelastic ageing and mechanoadaptation.

In conclusion, combining multiplexed trapping with harmonic decomposition could transform the method into a scalable mechanical imaging modality for living tissues. The ability to extract intrinsic viscoelastic spectra from intact, active and morphogenetic tissues could open up new avenues for studying mechanical feedback during development, quantifying in vivo tension remodelling, probing the mechanical signatures of differentiation or disease and linking nanoscale mechanics to tissue-scale morphodynamics.

Importantly, the method does not require optical transparency or mechanical isolation, making il applicable to complex trhee dimensional biological systems.

## 6 Conclusion

We have developed and validated a square-wave interferometric optical tweezers framework that is capable of quantitative microrheology in tissues with strong optical scattering and ATP-driven activity. By combining harmonic decomposition, global viscoelastic fitting and interferometric nanometric detection operating within the angular memory effect regime, we can measure cortical tension directly in diffusive millimetric tissues and extract viscoelastic properties at different frequencies under non-equilibrium, ATP-driven activity. The system is also robust to neighbouring cellular forces thanks to harmonic filtering. Unlike transmission-based microrheology approaches, our method does not rely on simplified optical propagation models, which are ineffective in heterogeneous tissues. Instead, it preserves quantitative force calibration and spectral reconstruction, even in the presence of strong multiple scattering and morphogenetic remodelling. This work establishes a link between interferometric optical physics and developmental tissue mechanics, providing a scalable, frequency-resolved mechanical probe for intact living systems. The development of this method is a breakthrough in mechanobiology, offering the ability to measure tension and stiffness with unprecedented precision in complex scattering tissues and in the presence of active noise.

## 7 Acknowledgments

We thank Marc Allain and Loïc Le Goff, Wylie Ahmed for support and fruitful discussions on the project.

This work was funded by the following agencies: Agence Nationale de la Recherche (ANR-20-CE45-0024,ANR-22-CE42-0026,ANR-21-CE13-0012);

## 8 Contributions

TME conducted the Drosophila leg and milk experiments, as well as the viscoelastic measurements. AP and NF designed the direct FDT method and the detector calibration without global fitting. RB performed the measurements on Drosophila embryos. VD performed nanodissection experiments on Drosophila pupae. APM designed the Drosophila pupae experiment. AS and TM designed the theoretical and experimental methods linked to the memory effect. TME, AP, NF, and TM designed the data analysis strategy. TME, AS, TM, and MS wrote the papers.

