## Supplementary Material for "Deep-tissue absolute force spectroscopy with sub-piconewton precision"

### 9 Supplementary Material

#### 9.1 The localization precision for QPD detectors or centroid-based detection is crucial.

For a QPD, the displacement signal along $x$ is obtained from the differential intensity between two pupil halves,

$$V_{\mathrm{QPD}} \propto \int_{k_x>0} I(\mathbf{k})\,\mathrm{d}\mathbf{k} - \int_{k_x<0} I(\mathbf{k})\,\mathrm{d}\mathbf{k}. \tag{18}$$

Linearizing for small displacements ($\Delta x \ll \lambda$) yields

$$V_{\mathrm{QPD}} \simeq C_{\mathrm{eff}}\, k_0 \mathrm{NA}\, \Delta x, \tag{19}$$

where $C_{\mathrm{eff}}$ is the effective interferometric contrast after speckle averaging.

The noise variance of the QPD signal results from the sum of speckle fluctuations over the pupil halves,

$$\sigma^2_{\mathrm{QPD}} \propto \frac{1}{N_{\mathrm{speckle}}}, \tag{20}$$

leading to a localization precision

$$\boxed{\delta x_{\mathrm{QPD}} \sim \frac{1}{k_0 \mathrm{NA}} \sqrt{N_{\mathrm{speckle}}}} \tag{21}$$

The key point is that the differential integration performed by the QPD averages out short-range speckle fluctuations, leaving only long-wavelength distortions to affect the cosine slope. In centroid detection, the displacement is estimated from the first moment of the intensity distribution,

$$x_c = \frac{\int x\, I(\mathbf{k})\,\mathrm{d}\mathbf{k}}{\int I(\mathbf{k})\,\mathrm{d}\mathbf{k}}. \tag{22}$$

Expanding $I(\mathbf{k})$ to first order in $\Delta x$ gives

$$x_c \simeq \frac{\int x\, |E_0(\mathbf{k})|^2 S(\mathbf{k}) \cos(k_x \Delta x)\,\mathrm{d}\mathbf{k}}{\int |E_0(\mathbf{k})|^2 S(\mathbf{k})\,\mathrm{d}\mathbf{k}}. \tag{23}$$

Because the centroid estimator explicitly weights the intensity by position, local speckle maxima contribute disproportionately to the signal. The noise variance therefore scales as

$$\sigma^2_{\mathrm{centroid}} \propto \frac{\langle x^2 \rangle}{N_{\mathrm{speckle}}}, \tag{24}$$

where $\langle x^2 \rangle$ is the second moment of the pupil distribution. As a consequence, the localization precision becomes

$$\boxed{\delta x_{\mathrm{centroid}} \sim \frac{\sigma_x}{k_0 \mathrm{NA}} \sqrt{N_{\mathrm{speckle}}}} \tag{25}$$

with $\sigma_x$ the characteristic pupil radius, yielding a significantly larger prefactor than in QPD detection.

#### 9.2 Custom-built setup dedicated to optical trapping measurements with QPD detector.

The system combines a conventional inverted fluorescence microscope with an active optical trapping module. The trapping system consists of two main components: a beam steering unit enabling precise positioning of the optical trap within the sample, and a back focal plane interferometry (BFPi) detection scheme used to track the position of the trapped object relative to the trap center.

Nanometric displacement of the optical trap is achieved using a steering mirror conjugated to the pupil plane of the microscope (Thorlabs FSM75-P01). A three-axis piezoelectric stage (PiezoConcept BIO3) is used to accurately position the sample and maintain focus on the trapped object. The setup is implemented on an inverted fluorescence microscope (Leica DMI6000 B) equipped with a 100× oil-immersion objective (NA = 1.4, Leica HCX PL APO). To optimize the filling of the objective back aperture by the infrared trapping beam (up to $\sim 85\%$), a ×4 afocal telescope composed of two relay lenses (Thorlabs ACA254-050-1064 and ACA254-200-1064) is used. The resulting diffraction-limited focal spot has a full width of approximately 600 nm, ensuring efficient optical trapping.

The system is configured for simultaneous fluorescence imaging (GFP and RFP) and infrared trapping using a multiband dichroic mirror (Semrock Di03-R405/488/561/635-t1-25x36). Synchronization between imaging and trapping is achieved using a National Instruments acquisition card combined with an Inscoper synchronization module, with all processes controlled via Inscoper software.

**Rapid focal sweep set-up:** the remote focusing system is based on active lenses in the collection part of the microscope. The raly lense are thorlabs AC254-125-A-ML,AC254-125-A-ML. The exact position of the ETL and its associated offset lens (Optotune EL-10-30-Ci Series) is finely adjusted so as to optimize the detection PSF. We found that placing specifically the ETL in the fourier plane and not the offset lens or the mid-point between the offset lens and the ETL was optimal. T Fluorescence is further filtered with a bandpass filter (Semrock, FF01-525/45-25 ). inscoper harware and software is done using though digital pulses and analog signal sent to the ETL driver.

#### 9.3 Commercial optical tweezers platform (SENSOCELL, Impetux)

Experiments were also performed using a commercial optical tweezers platform (SENSOCELL, Impetux) integrated on a Nikon inverted microscope (Tie2 series). The system combines a fluorescence imaging module with an advanced optical trapping unit coupled through the rear epifluorescence port of the microscope. Optical traps are generated using a 1064 nm infrared laser (maximum output power up to $\sim 5$ W) and focused through a high numerical aperture objective (typically a X60 water immersion NA 1.25), enabling efficient trapping in biological samples.

In contrast to the home-built setup, trap positioning is achieved using acousto-optic deflectors (AODs) conjugated to the objective pupil plane, allowing high-speed beam steering with update rates up to 25 kHz and the generation of multiple simultaneous traps (up to 256) within a field of view of approximately $80 \times 80$ $\mu$m . This configuration enables flexible control of trap trajectories, oscillations, and time-shared manipulation schemes.

Force and position measurements rely on a proprietary detection module based on light momentum conservation, which directly measures optical forces without requiring prior trap stiffness calibration [17]. This approach provides force sensitivity below 50 fN and typical position resolution on the order of 1 nm, with acquisition bandwidths up to [38].

#### 9.4 square wave signal, passive end active spectrum

The square-wave modulation of the trap position induces a periodic displacement of the trapped object. The differential BFPi signal can be written as

$$V(t) = \Re\left[\sum_{n=1,3,5,\ldots} -\beta^{-1}\frac{A}{n\pi}\frac{k_m(n\omega_d) - in\omega_d\gamma}{k_t + k_m(n\omega_d) - in\omega_d\gamma}\exp[-in\omega_d(t-t_0)]\right], \tag{26}$$

where $\beta$ is conversion factor of the BFPi system in $V.nm^{-1}$, A the amplitude of the square wave signal, $\omega_d$ the drive frequency, $k_m$ the viscoelastic response of the medium, $k_t$ the stiffness of the optical tweezers, $\gamma$ the friction coefficient of the medium, $t_0$ the dead time of the electronic acquisition device from the setup.

The fluctuation dissipation theorem (FDT) informs us that the power spectral density (PSD) link to thermal positional fluctuations (in $V2Hz^1$) is:

$$P_{\text{passive}}(\omega) = -\beta^{-2}\frac{2k_BT}{\omega}\Im\left[\frac{1}{k_m(\omega) + k_t + i\omega\gamma(\omega)}\right]. \tag{27}$$

The contribution to the PSD corresponding to the response of the bead to the laser drive is superimposed on this thermal spectrum. There are single data point spikes at all odd harmonics of the drive frequency when the bead displacement time series is truncated to an integer multiple of the drive period. The height of each peak is proportional to the measurement period as follow :

$$P_{active}(\omega) = \sum_{n=1,3,5...}^{\infty} -\beta^{-2}\frac{A^2 T_m}{\omega}[\frac{k_m n\omega_d - in\omega\gamma}{k_t + k_m n\omega_d - in\omega_d\gamma}]^2 \delta_{\omega,n\omega_d} \tag{28}$$

with $\delta_{\omega,n\omega_d}$ is 1 when $\omega = n\omega_d$ end zero elsewhere. An example of passive spectrum (red) and active spectrum (blue) is shown in figure 2.b between 0 and 500 Hz.

Each harmonic peak constitutes an independent measurement of the system response at frequency nd, enabling multiplexed calibration of the trap stiffness and detector sensitivity.

It is easy to estimate $\beta$ when a laser deflects rapidly, because it is possible to use a rectangular calibrated laser deflection with high accuracy, then $\delta X_{lasert=0} = \beta V_{t=0}$

#### 9.5 Multi-harmonic method without global fiting

A direct of the FDT calibration methode approach can be formulated by exploiting the multi-harmonic content of the square-wave excitation together with the independent calibration of the optical trap stiffness. Thanks to the active FDT calibration, the trap stiffness $k_t$ is first determined independently from the ratio of passive and active power spectral densities at each odd harmonic $n\omega_d$. Once $k_t$ is known, the mechanical response of the medium can be reconstructed independently at each harmonic frequency, without assuming a global rheological model. For each harmonic frequency $\omega = n\omega_d$, the complex susceptibility of the trapped bead is obtained from the passive fluctuations as:

$$\chi(\omega) = \frac{\langle |x(\omega)|^2 \rangle}{2k_B T} \tag{29}$$

The inverse susceptibility can then be written as:

$$\chi^{-1}(\omega) = k_t + 6\pi R\, G^*(\omega) \tag{30}$$

where $G^*(\omega) = G'(\omega) + iG''(\omega)$ is the effective complex viscoelastic modulus of the surrounding medium probed at frequency $\omega$ and $R$ is the radius of the trapped droplet. Thus, for each odd harmonic of the square-wave drive, the complex modulus is directly reconstructed as:

$$\boxed{G^*(n\omega_d) = \frac{1}{6\pi R}\left[\chi^{-1}(n\omega_d) - k_t\right]} \tag{31}$$

This multi-harmonic reconstruction provides independent estimates of $G'(\omega)$ and $G''(\omega)$ over a broad frequency range, without requiring any global fitting or prior assumption on the rheological model of the medium. The dispersion of the reconstructed values across harmonics further provides a direct estimate of experimental uncertainty and mechanical heterogeneity. When needed, a phenomenological viscoelastic model (e.g. power-law or Kelvin–Voigt-type behavior) can subsequently be fitted to $G'(\omega)$ and $G''(\omega)$ in a second step, without affecting the calibration of the optical trap.

#### 9.6 Global Fitting Method: Coupling the Estimation of Optical Tweezers Parameters and Mechanical Properties of Biological Media

Normally $k_t$, is constant for all frequencies. A mean value is taken from each harmonic to increase the accuracy of the tweezers constant spring.

In our study, the medium is considered like a cross linked polymer networks with the following equation based on [?] :

$$K_m(n\omega_d) = \sum_{n=1,3,5...}^{\infty} k_{m0}(n\omega_d) + k_{m1}(n\omega_d)\frac{iw^{\alpha}}{(\alpha-1)!} \tag{32}$$

with $\alpha$ the damping coefficient. Close to zero the medium is very elastic, close to 1 it's a fluid fluid. The elasticity component $k_{m0}$ is necessary when the droplet is in contact on elastic component like cell cortex or membrane.

After initialization of measured parameter $\beta$ and $k_t$, we can estimate more precisely all parameters $\beta$, $k_t$, $k_{m0}(n\omega_d)$, $k_{m1}(n\omega_d)$, $\gamma$ and $\alpha$ by minimizing simultaneously the functions $P_{passive}(\omega)$ and $P_{active}(\omega)$

$$\begin{array}{c} k_{m0}(n\omega_d), k_{m1}(n\omega_d), \\ \gamma, \alpha, \\ \beta, k_t \end{array} = ArgMin(P_{passive}(\omega), P_{active}(\omega)) \left\{ \begin{array}{c} 15pN \leqslant k_t \leqslant 35pN \\ 0 \leqslant k_{m0} \leqslant 20pN \\ 0 \leqslant k_{m1} \leqslant 20pN \\ 0 \leqslant \alpha \leqslant 1 \\ 10-8kg.s^-1 \leqslant \gamma \leqslant 10-7kg.s-1 \end{array} \right. \tag{33}$$

Note that, thanks to the knowledge of $\gamma$, in the case of a fluid, the viscosity can be estimated if the radius of the droplet is known.

$$n_{(n\omega)} = \frac{\gamma_{(n\omega)}}{6\pi R} \tag{34}$$

in the case of complex medium from $k_{m0}(n\omega_d)$, $k_{m1}(n\omega_d)$ and $\gamma$ the viscoelastic modulus of the medium $G(\omega)$ can be known

$$G(\omega) = G'(\omega) + iG''(\omega) \tag{35}$$

$$G(\omega) = \frac{1}{6\pi R}(k_{m0}(n\omega_d) + k_{m1}(n\omega_d)\frac{iw^\alpha}{(\alpha-1)!} + i\omega\gamma_{(n)}) \tag{36}$$

##### 9.7 Precision of the optical trap stiffness calibration in turbid media

The relative uncertainty on a single-harmonic estimate of $k_t$ is primarily limited by thermal noise, detector noise, and the finite acquisition time $T_m$, leading to

$$\frac{\delta k_t}{k_t} \sim \frac{1}{\sqrt{T_m \Delta f}}, \tag{37}$$

where $\Delta f$ is the effective spectral resolution. For typical experimental conditions ($T_m \approx 17$ s), this corresponds to a relative uncertainty of approximately 5–10% per harmonic. Crucially, the square-wave excitation enables simultaneous calibration over multiple harmonics. By averaging over $N_{\text{harm}}$ independent odd harmonics (typically 15–25 in our experiments, up to $\sim$ 500 Hz), the statistical uncertainty on the trap stiffness is reduced as

$$\frac{\delta k_t}{k_t} \sim \frac{1}{\sqrt{N_{\text{harm}}}} \left(\frac{\delta k_t}{k_t}\right)_{\text{single}}, \tag{38}$$

yielding a final relative precision of approximately 1–3%. For typical trap stiffness values in this study ($k_t \sim$ 15–35 pN/$\mu$m), this corresponds to an absolute uncertainty of

$$\delta k_t \sim 0.3\text{–}1 \text{ pN}/\mu\text{m}, \tag{39}$$

even in strongly scattering biological tissues. Importantly, optical diffusion primarily degrades the instantaneous signal-to-noise ratio but does not bias the stiffness estimation, as the calibration relies on spectral ratios and is internally renormalized at each measurement.

In living cells and tissues, the ultimate precision of force and rheological measurements using optical tweezers is not only limited by instantaneous localization noise, but also by stochastic intracellular activity and optical scattering. Importantly, when the optical trap is driven by a periodic square-wave displacement, the mechanical response is phase-locked to the drive, enabling efficient temporal averaging over multiple cycles.

For a square-wave displacement of amplitude $A$ and fundamental frequency $f_0$, the bead displacement contains odd harmonics at frequencies $nf_0$ with amplitudes

$$X_n = \frac{4A}{\pi n}, \quad n = 1, 3, 5, \ldots \tag{40}$$

Let $\sigma_x$ denote the instantaneous localization precision of the back focal plane interferometry (BFPi) system, which includes contributions from shot noise, speckle-induced contrast loss, and detector noise. When acquiring data over a total duration $T$, each harmonic component can be averaged over approximately $N = Tf_0$ independent cycles. The effective uncertainty on the amplitude of the $n$-th harmonic therefore scales as

$$\sigma_{x,n}^{\mathrm{eff}} \simeq \frac{\sigma_x}{\sqrt{Tf_0}}. \tag{41}$$

A harmonic is considered experimentally accessible when its amplitude exceeds the effective noise level,

$$\frac{4A}{\pi n} \gtrsim \frac{\sigma_x}{\sqrt{Tf_0}}. \tag{42}$$

This condition defines a maximum observable harmonic order,

$$n_{\max} \simeq \frac{4A}{\pi\sigma_x}\sqrt{Tf_0}, \tag{43}$$

and therefore a maximum accessible frequency,

$$\boxed{f_{\max} \simeq f_0\,\frac{4A}{\pi\sigma_x}\sqrt{Tf_0}.} \tag{44}$$

This expression highlights three key features of square-wave FDT calibration in turbid living media: (i) increasing the displacement amplitude $A$ linearly extends the accessible frequency range; (ii) improving localization precision $\sigma_x$ directly enhances high-frequency sensitivity; (iii) most importantly, the accessible frequency range increases as $\sqrt{T}$ with measurement time, making long acquisitions particularly powerful for suppressing stochastic intracellular noise such as ATP-driven activity.

In practice, this temporal averaging allows reliable extraction of trap stiffness and viscoelastic moduli at frequencies well above the fundamental drive frequency, even in strongly scattering tissues. This behavior fundamentally distinguishes square-wave multiplexed FDT calibration from single-frequency sinusoidal approaches, for which high-frequency sensitivity rapidly degrades in noisy biological environments.

#### 9.8 Precision on $G'$ and $G''$ limited by optical localization

The precision on the viscoelastic moduli extracted from fluctuation–dissipation optical tweezers measurements is ultimately limited by the spatial localization accuracy of the trapped probe. In complex and turbid media, the back focal plane interferometry (BFPI) signal arises from a noisy interference pattern, whose contrast is reduced by speckle mixing. This effect directly degrades the precision on the bead displacement $x(t)$ and therefore propagates to the estimation of $G'(\omega)$ and $G''(\omega)$.

For a spherical probe of radius $R$, the complex shear modulus is obtained as

$$G^*(\omega) = G'(\omega) + iG''(\omega) = \frac{1}{6\pi R}\left[k_m(\omega) + i\omega\gamma(\omega)\right]. \tag{45}$$

The uncertainty on $G^*$ follows directly from the uncertainty on the bead position $\delta x$, which affects both the passive power spectral density and the active response function. For a driven measurement at angular frequency $\omega$, the displacement amplitude is estimated as

$$x(\omega) = \frac{F(\omega)}{k_{\mathrm{trap}} + k_m(\omega) + i\omega\gamma}. \tag{46}$$

A localization uncertainty $\delta x$ leads to an uncertainty on the response function $\chi(\omega)$, and therefore on $k_m(\omega)$ and $\gamma(\omega)$. To first order, the relative error on the complex modulus reads

$$\frac{\delta G^*(\omega)}{|G^*(\omega)|} \simeq \frac{\delta x}{x(\omega)}. \tag{47}$$

Separating elastic and viscous contributions yields

$$\delta G'(\omega) \simeq \frac{k_m(\omega)}{6\pi R}\frac{\delta x}{x(\omega)}, \tag{48}$$

$$\delta G''(\omega) \simeq \frac{\omega\gamma(\omega)}{6\pi R}\frac{\delta x}{x(\omega)}. \tag{49}$$

Thus, the precision on $G'$ and $G''$ scales linearly with the localization noise and inversely with the driven displacement amplitude. The localization precision in the back focal plane depends on the numerical aperture (NA) of the detection optics and on the effective contrast of the interference signal. For a trapped bead of diameter 1 $\mu$m, typical values are

- NA = 0.55 (home-made system): $\delta x \simeq 20$–$40$ nm in turbid tissue,
- NA = 1.4 (commercial system): $\delta x \simeq 5$–$15$ nm under similar scattering conditions.

Since $G^*$ scales as $1/R$, larger probes reduce the relative uncertainty on the mechanical moduli. For $R = 0.5$ $\mu$m, the conversion factor is

$$\frac{1}{6\pi R} \simeq 1.06 \times 10^5 \ \mathrm{Pa\,m^{-1}}. \tag{50}$$

For a driven displacement amplitude of $x(\omega) = 100$ nm, the expected precision is:

- $\delta x = 5$ nm $\Rightarrow \delta G'/G' \sim 5\%$,
- $\delta x = 20$ nm $\Rightarrow \delta G'/G' \sim 20\%$,
- $\delta x = 40$ nm $\Rightarrow \delta G'/G' \sim 40\%$.

In absolute terms, for a typical cellular modulus $G' \sim 100$–$500$ Pa, this corresponds to

$$\delta G' \sim 5\text{–}25 \text{ Pa} \quad (\text{NA} = 1.4), \qquad \delta G' \sim 20\text{–}200 \text{ Pa} \quad (\text{NA} = 0.55). \tag{51}$$

The viscous modulus $G''$ is generally more sensitive to localization noise at high frequency due to the $\omega$ prefactor, which ultimately limits the highest reliable harmonic accessible in square-wave or sinusoidal driving experiments. These results highlight that the achievable precision on $G'$ and $G''$ in turbid biological media is primarily limited by the contrast of the back focal plane interference pattern. Increasing numerical aperture, probe size, and driving amplitude improves precision, whereas an increasing number of speckle modes reduces it. This trade-off defines the optimal experimental window for high-frequency microrheology in living tissues.

##### 9.9 Estimation of uncertainties on G' and G" via statistical averaging and frequency

In our optical measurements, the localization precision $\sigma_x$ directly limits the estimation of viscoelastic moduli. When a square-wave displacement is applied, the effective number of independent realizations per frequency, denoted $N_m(\omega)$, is proportional to the number of cycles measured:

$$N_m(\omega) = f \cdot T_{\text{measure}}, \tag{52}$$

where $f$ is the frequency of the applied signal and $T_{\text{measure}}$ is the total measurement duration. The statistical reduction of phase uncertainty $\sigma_\phi$ then follows:

$$\sigma_\phi(\omega) = \frac{\sigma_x}{A\sqrt{N_m(\omega)}} = \frac{\sigma_x}{A\sqrt{f T_{\text{measure}}}}, \tag{53}$$

with $A$ being the displacement amplitude imposed by the trap. Thus, for a 200 nm square-wave at 10 Hz over 15 s, we obtain $N_m = 150$ and a relative phase uncertainty of $\sigma_\phi \approx 0.41\%$.

The propagation of this uncertainty onto the viscoelastic moduli can be expressed as:

$$\delta G' \sim G' \cdot \sigma_\phi + G_0 \cdot \frac{\sigma_x}{A} \cdot \text{amplitude factor}, \qquad \delta G'' \sim G'' \cdot \sigma_\phi + G_0 \cdot \frac{\sigma_x}{A} \cdot \text{amplitude factor}, \tag{54}$$

where the second term corresponds to additive detection noise, which is particularly important for G' at high frequencies. The amplitude factor can be adjusted to match the experimentally observed noise level. Therefore, even though $N_m$ increases with $f$, the relative noise on G" rises at high frequencies, while G' remains less noisy but receives a realistic correction from additive detection noise.

This approach correctly captures:

- the frequency dependence of statistical noise,
- the difference in noise between G' and G",
- and the reduction of uncertainty by increasing the measurement duration $T_{\text{measure}}$.

For a 15 s measurement with a 200 nm square-wave at 10 Hz, $N_m = 150$, and the phase is known to $\sim 0.41$ %, resulting in realistic relative uncertainties on G' and G" compatible with experimental observations.

#### 9.10 theoritical model of the Milk

To accurately describe the viscoelastic response measured in our experiments, we employed an extended Maxwell model incorporating both a residual elastic contribution and a weak power-law component. The complex shear modulus is expressed as

$$G^*(\omega) = G_\infty + \frac{G_0\, i\omega\tau}{1 + i\omega\tau} + G_1(i\omega)^\alpha,$$

where $G_\infty$ accounts for a low-frequency elastic contribution, $G_0$ and $\tau$ define the characteristic Maxwell relaxation, and the term $G_1(i\omega)^\alpha$ captures a broad distribution of relaxation processes typically observed in soft disordered materials. This additional component provides a better description of the intermediate frequency regime, where deviations from a single-mode Maxwell behavior are consistently observed.

To reproduce experimental conditions, we further incorporate a noise model arising from finite localization precision and active fluctuations. The relative uncertainty on the measured moduli is described as

$$\frac{\delta G}{G} \sim \frac{\sigma_x}{A\sqrt{N_m}} \sqrt{1 + \left(\frac{\omega}{\omega_c}\right)^2},$$

where $\sigma_x$ is the localization precision, $A$ the drive amplitude, $N_m$ the number of statistically independent measurements, and $\omega_c$ the characteristic crossover frequency of the system. This formulation captures both the increase of uncertainty at high frequency due to reduced signal amplitude and the contribution of active fluctuations at low frequency.

Such extended viscoelastic descriptions are consistent with previous studies on complex fluids and biological materials, including casein-based emulsions and gels, which exhibit Maxwell-like behavior combined with power-law rheology over broad frequency ranges. In particular, dairy systems such as casein micelle suspensions and acid-induced gels have been shown to display a dominant relaxation mode together with distributed relaxation spectra arising from microstructural heterogeneity and interparticle interactions. These features have been extensively characterized in microrheology and bulk rheology studies (e.g. Lucey et al., Int. Dairy Journal, 2003; Horne, Curr. Opin. Colloid Interface Sci., 2002; Dalgleish, J. Dairy Sci., 2011), and provide a relevant physical framework for interpreting the mechanical response observed in our experiments.

#### 9.11 Minimum Time Window Required for Reliable Determination of $G''(\omega)$ up to 300 Hz

The determination of the loss modulus $G''(\omega)$ from harmonic microrheology critically depends on the accuracy of phase estimation between the driven and response signals. The relative error on $G''$ is dominated by the uncertainty in the measured phase $\phi(\omega)$ and can be approximated as

$$\frac{\delta G''}{G''} \simeq \frac{\delta\phi}{\tan\phi}. \tag{55}$$

This relation highlights that $G''$ becomes ill-conditioned when $\phi \to 0$ (elastic-dominated regime) or $\phi \to \pi/2$ (viscous-dominated regime). In practice, to maintain a relative error below 10%, one requires

$$\tan\phi > 0.17 \quad \Rightarrow \quad 10^\circ \lesssim \phi \lesssim 80^\circ. \tag{56}$$

The phase uncertainty at harmonic frequency $f_n = nf_d$ scales as

$$\delta\phi \sim \frac{1}{\sqrt{Tf_n \cdot \mathrm{SNR}}}, \tag{57}$$

where $T$ is the acquisition duration and SNR is the signal-to-noise ratio at the considered harmonic. Since the number of oscillation cycles contributing to the estimate is $N_n = Tf_n$, phase precision improves as $\sqrt{Tf_n}$.

Therefore, to reliably extract $G''(\omega)$ up to a target frequency $f_{\max}$, the time window must satisfy

$$T \geq \frac{1}{f_{\max}} \left( \frac{1}{\delta\phi_{\text{target}}^2 \cdot \mathrm{SNR}} \right). \tag{58}$$

Assuming a conservative phase precision target $\delta\phi_{\text{target}} \approx 1^\circ = 0.017$ rad and SNR $\sim 10$, one obtains

$$Tf_{\max} \gtrsim 350, \tag{59}$$

which provides a practical design criterion.

**Case 1: 1 kHz sampling, $f_d = 2.49$ Hz.**

To avoid spectral leakage, the acquisition time must contain an integer number of cycles:

$$T = \frac{N}{f_d}. \tag{60}$$

Choosing $N = 17$ cycles yields $T = 6.83$ s. The frequency resolution is then $\Delta f = 1/T = 0.146$ Hz, sufficient to isolate harmonics up to 300 Hz. At $f_{\max} = 300$ Hz, the number of cycles contributing to phase estimation is

$$N_{300} = Tf_{\max} \approx 2050, \tag{61}$$

ensuring $\delta\phi < 1^\circ$ and therefore stable determination of $G''$. Durations below $\sim 5$ s significantly degrade low-frequency phase precision and compromise the conditioning of $G''$.

**Case 2: 1 kHz sampling, $f_d = 1.42$ Hz.**

Imposing integer cycle acquisition, selecting $N = 24$ cycles gives

$$T = 16.90 \text{ s}. \tag{62}$$

The corresponding resolution is $\Delta f = 0.059$ Hz. At 300 Hz,

$$N_{300} = Tf_{\max} \approx 5070, \tag{63}$$

providing excellent phase precision across the full spectral window. Reducing the acquisition to $T \sim 10$ s (14 cycles) remains acceptable but increases $\delta\phi$ at low harmonics and narrows the reliable $G''$ range.

**Case 3: 25 kHz sampling, $f_d = 1$ Hz.**

With $T = 10$ s (10 cycles), $\Delta f = 0.1$ Hz. At 300 Hz,

$$N_{300} = 3000, \tag{64}$$

which ensures phase errors well below 1°. Increasing the duration to 20 s further improves conditioning but yields diminishing returns beyond $Tf_{\max} \sim 3000$.

**Minimum Practical Time Window.**

For square-wave harmonic microrheology targeting reliable extraction of $G''(\omega)$ up to 300 Hz, a robust empirical criterion is

$$T \gtrsim \frac{15}{f_d}, \tag{65}$$

corresponding to at least 15 fundamental cycles. This guarantees:

- negligible spectral leakage (integer cycle acquisition),
- phase uncertainty below $\sim 1^\circ$ at high harmonics,
- stable conditioning of $G''$ within the physically informative phase window.

In all considered configurations, the mechanical bandwidth (not the electronic sampling rate) sets the ultimate limit. The acquisition duration must therefore be chosen to optimize phase precision rather than frequency resolution alone.

#### 9.12 Resolution on membrane tension and elastic modulus under active fluctuation in drosophila embryo

For a measurement duration $T_m$ spanning several drive periods, the uncertainty on the complex displacement amplitude at harmonic $n$ scales as

$$\delta x_n \simeq \sqrt{\frac{S_{\text{act}}(n\omega_d)}{T_m}}, \tag{66}$$

where $S_{\text{act}}(\omega) \sim \omega^{-\alpha}$ is the active fluctuation spectrum. As a result, higher harmonics benefit from both spectral separation from low-frequency activity and reduced active noise (sea section 4.3).

Experimentally, for typical acquisition times ($T_m \sim 15$–$30$ s), we estimate:

$$\delta x_{n=1} \approx 15\text{–}20 \text{ nm}, \qquad \delta x_{n=3} \approx 8\text{–}10 \text{ nm}, \qquad \delta x_{n=5} \approx 5\text{–}7 \text{ nm}.$$

The force resolution at each harmonic follows directly from the calibrated trap stiffness:

$$\delta F_n = k_t \, \delta x_n. \tag{67}$$

Using $k_t \simeq 75$ pN/$\mu$m, this yields:

$$\delta F_{n=1} \approx 1.1\text{–}1.5 \text{ pN}, \quad \delta F_{n=3} \approx 0.6\text{–}0.8 \text{ pN}, \quad \delta F_{n=5} \approx 0.4\text{–}0.5 \text{ pN}.$$

In the quasi-elastic regime relevant for nuclear membrane deformation at these frequencies, the effective membrane tension $\sigma$ is inferred from the plateau force response. Given the imposed displacement amplitude $\Delta x = 200$ nm, the relative uncertainty on the extracted tension is therefore

$$\frac{\delta\sigma}{\sigma} \simeq \frac{\delta x_n}{\Delta x}, \tag{68}$$

leading to relative precisions of approximately 10%, 5%, and 3% at the first, third, and fifth harmonics, respectively.

Within the linear response framework, the storage modulus is obtained from the in-phase component of the complex force–displacement ratio:

$$G'(n\omega_d) \propto \text{Re}\left[\frac{F_n}{x_n}\right]. \tag{69}$$

Error propagation yields

$$\frac{\delta G'}{G'} \simeq \sqrt{\left(\frac{\delta F_n}{F_n}\right)^2 + \left(\frac{\delta x_n}{x_n}\right)^2}. \tag{70}$$

Because the imposed displacement amplitude (200 nm) is much larger than the residual uncertainty $\delta x_n$, the dominant contribution arises from displacement noise. We therefore estimate a relative precision on $G'$ of order:

$$\delta G'/G' \approx 10\% \text{ at } n=1, \quad 5\% \text{ at } n=3, \quad \text{and } \lesssim 4\% \text{ at } n=5.$$

Importantly, independent slow square-step relaxation experiments performed at lower frequencies yield elastic plateaus consistent with the values of $G'$ extracted from the higher-frequency square-wave harmonics. This agreement confirms that, despite strong ATP-driven stochastic activity, the elastic component of the nuclear membrane response is reliably captured by small-amplitude, higher-frequency square-wave probing.

Overall, these results demonstrate that sub-piconewton force sensitivity and few-percent accuracy on membrane elasticity can be achieved in living embryos, even in the presence of large active fluctuations, provided that measurements are performed in the frequency domain and rely on phase-locked detection.

#### 9.13 Resolution on membrane tension and elastic modulus under active fluctuation in drosophila embryo

For a measurement duration $T_m$ spanning several drive periods, the uncertainty on the complex displacement amplitude at harmonic $n$ scales as

$$\delta x_n \simeq \sqrt{\frac{S_{\text{act}}(n\omega_d)}{T_m}}, \tag{71}$$

where $S_{\text{act}}(\omega) \sim \omega^{-\alpha}$ is the active fluctuation spectrum. As a result, higher harmonics benefit from both spectral separation from low-frequency activity and reduced active noise (sea section 4.3).

Experimentally, for typical acquisition times ($T_m \sim$ 15–30 s), we estimate:

$$\delta x_{n=1} \approx 15\text{–}20 \text{ nm}, \qquad \delta x_{n=3} \approx 8\text{–}10 \text{ nm}, \qquad \delta x_{n=5} \approx 5\text{–}7 \text{ nm}.$$

The force resolution at each harmonic follows directly from the calibrated trap stiffness:

$$\delta F_n = k_t \, \delta x_n. \tag{72}$$

Using $k_t \simeq 75$ pN/$\mu$m, this yields:

$$\delta F_{n=1} \approx 1.1\text{–}1.5 \text{ pN}, \quad \delta F_{n=3} \approx 0.6\text{–}0.8 \text{ pN}, \quad \delta F_{n=5} \approx 0.4\text{–}0.5 \text{ pN}.$$

In the quasi-elastic regime relevant for nuclear membrane deformation at these frequencies, the effective membrane tension $\sigma$ is inferred from the plateau force response. Given the imposed displacement amplitude $\Delta x = 200$ nm, the relative uncertainty on the extracted tension is therefore

$$\frac{\delta \sigma}{\sigma} \simeq \frac{\delta x_n}{\Delta x}, \tag{73}$$

leading to relative precisions of approximately 10%, 5%, and 3% at the first, third, and fifth harmonics, respectively.

Within the linear response framework, the storage modulus is obtained from the in-phase component of the complex force–displacement ratio:

$$G'(n\omega_d) \propto \text{Re}\left[\frac{F_n}{x_n}\right]. \tag{74}$$

Error propagation yields

$$\frac{\delta G'}{G'} \simeq \sqrt{\left(\frac{\delta F_n}{F_n}\right)^2 + \left(\frac{\delta x_n}{x_n}\right)^2}. \tag{75}$$

Because the imposed displacement amplitude (200 nm) is much larger than the residual uncertainty $\delta x_n$, the dominant contribution arises from displacement noise. We therefore estimate a relative precision on $G'$ of order:

$$\delta G'/G' \approx 10\% \text{ at } n = 1, \quad 5\% \text{ at } n = 3, \quad \text{and } \lesssim 4\% \text{ at } n = 5.$$

Importantly, independent slow square-step relaxation experiments performed at lower frequencies yield elastic plateaus consistent with the values of $G'$ extracted from the higher-frequency square-wave harmonics. This agreement confirms that, despite strong ATP-driven stochastic activity, the elastic component of the nuclear membrane response is reliably captured by small-amplitude, higher-frequency square-wave probing.

Overall, these results demonstrate that sub-piconewton force sensitivity and few-percent accuracy on membrane elasticity can be achieved in living embryos, even in the presence of large active fluctuations, provided that measurements are performed in the frequency domain and rely on phase-locked detection.

#### 9.14 Sub piconewton precision of nuclear membrane tension in drosophila embryo in presence of highly ATP dependent process

For a trapped probe embedded in a living cell or tissue, the measured displacement along one spatial direction can be decomposed as

$$x(t) = x_{\rm trap}(t) + x_{\rm th}(t) + x_{\rm act}(t) + x_{\rm ext}(t), \tag{76}$$

where $x_{\rm trap}$ denotes the deterministic response to the optical trap, $x_{\rm th}$ the thermal fluctuations governed by the fluctuation–dissipation theorem, $x_{\rm act}$ the active intracellular fluctuations driven by ATP-dependent processes, and $x_{\rm ext}$ the contribution arising from long-range mechanical coupling to neighboring cells.

In living systems, active fluctuations dominate thermal noise over a broad frequency range, particularly at low frequencies, and therefore impose a fundamental limit to localization precision.

Experimental and theoretical studies have shown that active intracellular forces exhibit a power-law spectrum of the form

$$S_{\rm act}(\omega) \sim \frac{A_{\rm act}}{\omega^{\alpha}}, \qquad 1 < \alpha < 2, \tag{77}$$

reflecting the non-equilibrium nature of force generation by molecular motors and cytoskeletal remodeling. As a consequence, the fluctuation–dissipation theorem is violated at low frequencies, and the variance of particle position is no longer bounded by thermal noise alone.

The total position variance can be expressed as

$$\sigma_x^2 = \int_0^{\omega_c} |\chi(\omega)|^2 \left[S_{\rm th}(\omega) + S_{\rm act}(\omega)\right] d\omega + \sigma_{\rm det}^2, \tag{78}$$

where $\chi(\omega)$ is the mechanical susceptibility of the trapped particle and $\sigma_{\rm det}$ is the detection-limited precision discussed in the previous section.

Even with nanometric optical tracking capabilities, the effective precision in metabolically active cells typically degrades to the 10–30 nm range due to active noise. the Fig 3.A typically shown the active fluctuation of the embryo drosophila

#### 9.15 Spatial locality of mechanical measurements from square-wave harmonics in drosophila leg

In addition to optical and detection limitations, intracellular and intercellular mechanical coupling introduces long-range spatial correlations in membrane displacements. Experimentally, we observe displacement correlations extending up to $\sim 10\ \mu$m in wild-type tissues and $\sim 2\ \mu$m in mechanically altered mutants, consistent with a Gaussian spatial correlation function,

$$C(r,\omega) = \langle x(0,\omega)x(r,\omega)\rangle = C_0(\omega)\exp\left(-\frac{r^2}{2\xi^2(\omega)}\right), \tag{79}$$

where $\xi(\omega)$ denotes the frequency-dependent mechanical correlation length.

In viscoelastic media, $\xi(\omega)$ decreases with frequency due to limited stress propagation, and is typically described by a diffusive scaling law,

$$\xi(\omega) \propto \omega^{-1/2}. \tag{80}$$

Consequently, higher-frequency mechanical perturbations probe increasingly local material properties.

Square-wave excitation naturally provides access to a set of odd harmonics $\omega_n = n\omega_d$ ($n = 1, 3, 5, \dots$), each probing a characteristic correlation length

$$\xi_n = \xi(\omega_n) = \xi(\omega_d)/\sqrt{n}. \tag{81}$$

For example, for a correlation length $\xi(\omega_d) = 10$ $\mu$m at the fundamental frequency, the third and fifth harmonics probe length scales of approximately 5.8 $\mu$m and 4.5 $\mu$m, respectively. In mechanically altered mutants with $\xi(\omega_d) \sim 2$ $\mu$m, the third harmonic already probes sub-micrometer length scales.

Therefore, the extraction of the storage modulus $G'(\omega)$ at higher harmonics provides increasingly local mechanical information, even in the presence of long-range intercellular mechanical coupling. In our experiments, the combination of square-wave amplitudes of 100–300 nm, trap stiffness $k_t \sim 130$ pN/$\mu$m, and numerical apertures of 0.55 and 1.4 ensures that optical localization precision remains below 20 nm, such that the dominant limitation to locality arises from biological mechanical correlations rather than optical noise.

#### 9.16 Local and non-local mechanical response

Mechanical measurements performed on a single cell embedded in living tissue are influenced by the transmission of force through the surrounding cellular network. To disentangle the effects of intrinsic cellular mechanics from those of the tissue as a whole, we model the measured displacement as the sum of a local response and a non-local contribution arising from force propagation.

We denote by $x_0(t)$ the displacement of the cell membrane or cytosol along the trapping axis. In Fourier space, the measured response reads

$$x_0(\omega) = \chi_{\text{eff}}(\omega)\, F_{\text{opt}}(\omega), \tag{82}$$

where $F_{\text{opt}}(\omega)$ is the optically applied force and $\chi_{\text{eff}}(\omega)$ the effective mechanical susceptibility.

The optical force is generated by controlled displacements of the laser focus :

$$F_{\text{opt}}(\omega) = k_{\text{trap}}\big[x_L(\omega) - x_0(\omega)\big], \tag{83}$$

with $k_{\text{trap}}$ the trap stiffness correctly calibrated and $x_L(t)$ the imposed laser displacement

The effective susceptibility is decomposed as

$$\chi_{\text{eff}}^{-1}(\omega) = k_{\text{trap}} + i\omega\big[\gamma_{\text{cell}}(\omega) + \gamma_{\text{tissue}}(\omega)\big], \tag{84}$$

where $\gamma_{\text{cell}}(\omega)$ describes the intrinsic viscoelastic dissipation of the probed cell, and $\gamma_{\text{tissue}}(\omega)$ accounts for mechanical coupling to the surrounding tissue.

To quantify force propagation within the tissue, we simultaneously track the displacement of neighboring cell vertices. The displacement field $u(r, \omega)$ at a distance $r$ from the trapped cell is modeled as

$$u(r, \omega) = G_m(r, \omega)\, F_{\text{opt}}(\omega), \tag{85}$$

where $G_m(r, \omega)$ is an effective mecanical Green function of the tissue (see apendix)

By fitting the spatial decay of vertex displacements, we extract $\xi(\omega)$ and identify a frequency regime in which $\xi(\omega)$ becomes smaller than the characteristic cell size. In this regime, non-local mechanical contributions are strongly attenuated and the measured response is dominated by intrinsic cellular mechanics.

#### 9.17 Effective mechanical Green function of a viscoelastic tissue

We model the tissue as an effective isotropic viscoelastic continuum characterized by a complex shear modulus $G^*(\omega)$. In the overdamped regime, the displacement field $\mathbf{u}(\mathbf{r}, \omega)$ induced by a localized force $\mathbf{F}(\omega)$ obeys

$$-\nabla \cdot \boldsymbol{\sigma}(\mathbf{r}, \omega) + \eta_{\text{eff}}(\omega)\, i\omega \mathbf{u}(\mathbf{r}, \omega) = \mathbf{F}(\omega)\delta(\mathbf{r}), \tag{86}$$

where $\boldsymbol{\sigma}$ is the stress tensor and $\eta_{\text{eff}}(\omega)$ an effective frequency-dependent viscosity.

The corresponding Green function relates force and displacement as

$$u_i(\mathbf{r}, \omega) = G_{ij}(\mathbf{r}, \omega)\, F_j(\omega). \tag{87}$$

In three dimensions, the scalar part of the Green function reads

$$G(r,\omega) = \frac{1}{4\pi G^*(\omega) r} \exp\left(-\frac{r}{\xi(\omega)}\right), \tag{88}$$

with the mechanical correlation length

$$\xi(\omega) = \sqrt{\frac{G^*(\omega)}{i\omega\eta_{\text{eff}}(\omega)}}. \tag{89}$$

This form captures the crossover between long-range elastic force propagation at low frequencies and localized, dissipative mechanical response at high frequencies. The experimentally measured decay of vertex displacements provides a direct estimate of $\xi(\omega)$ and allows us to identify the frequency regime in which single-cell mechanical properties can be reliably extracted.

#### 9.18 Correction of optical moment measurements in turbid tissues for commercial systems

In commercial optical tweezers systems, force measurements are often obtained by direct detection of the optical momentum transfer in the back focal plane of a high numerical aperture condenser. In homogeneous and weakly scattering media, this approach provides an absolute force measurement without requiring prior calibration of the position detector or trap stiffness. However, in optically turbid biological tissues, multiple scattering strongly alters the angular distribution of the collected light, leading to systematic errors in the measured optical momentum. To investigate whether square-wave laser driving could be used to correct optical momentum measurements in scattering media, we consider a transverse square-wave displacement of the optical trap,

$$x_{\text{trap}}(t) = A\,\text{sgn}[\sin(\omega_d t)], \tag{90}$$

where $A$ is the displacement amplitude and $\omega_d$ the drive angular frequency. The corresponding Fourier decomposition contains only odd harmonics,

$$x_{\text{trap}}(t) = \sum_{n=1,3,5\ldots}^{\infty} \frac{4A}{n\pi} \sin(n\omega_d t). \tag{91}$$

In the linear regime of the optical trap, the true force applied to the trapped object at each harmonic is given by

$$F_n^{\text{true}} = k_t \frac{4A}{n\pi}, \tag{92}$$

where $k_t$ is the optical trap stiffness. In an ideal momentum-detection system, the measured optical momentum signal $M_n$ would be directly proportional to $F_n^{\text{true}}$. In the presence of a turbid medium, however, the measured optical momentum is modified by an unknown, frequency-dependent transfer function $\eta_n$,

$$M_n^{\text{meas}} = \eta_n F_n^{\text{true}}. \tag{93}$$

The factor $\eta_n$ accounts for multiple scattering, reduced effective numerical aperture, and distortion of the angular light distribution in the pupil plane. Importantly, $\eta_n$ depends on the local optical properties of the tissue, the depth of the trap, and the spatial configuration of the scattering structures, and therefore cannot be assumed to be constant or known *a priori*. In principle, one could attempt to correct the optical momentum measurement by estimating $\eta_n$ from the known square-wave displacement amplitude,

$$\eta_n = \frac{M_n^{\text{meas}}}{k_t\, 4A/(n\pi)}. \tag{94}$$

However, this approach requires independent knowledge of the trap stiffness $k_t$, which is itself altered by the viscoelastic and heterogeneous nature of living tissues. As a consequence, the calibration problem becomes underdetermined: both $k_t$ and $\eta_n$ are unknown and cannot be disentangled from momentum measurements alone. Moreover, unlike position-based calibration methods relying on the fluctuation–dissipation theorem, optical momentum detection lacks a direct thermodynamic constraint linking the driven response to thermal fluctuations. This absence of an Onsager-type reciprocity prevents self-consistent calibration of the force detector in complex biological environments.

As a result, although square-wave driving introduces well-defined harmonic components in the optical momentum signal, it cannot provide an absolute and robust force calibration in turbid media. The inferred forces remain renormalized by an unknown, sample-dependent factor, leading to systematic errors that cannot be corrected without additional assumptions.

By contrast, fluctuation–dissipation-based calibration combined with back focal plane interferometry enables simultaneous determination of the position detector sensitivity, trap stiffness, and viscoelastic response of the surrounding medium. This self-consistent approach yields absolute force measurements with sub-piconewton precision, even in strongly scattering living tissues

#### 9.19 Laser nano dissection

Laser ablation experiments were performed using a pulsed, Q-switched Nd:YAG laser (532 nm wavelength, 0.4 ns pulse duration, up to 7 kHz repetition rate and 7 J energy per pulse) coupled to an ILAS2 galvanometer-based scanning system (Roper Scientific), which was mounted on an inverted Leica DMI6000B microscope. The laser beam was focused through a high-numerical-aperture oil-immersion objective (Plan-Apochromat ×100/0.7–1.4, Leica).

Cell–cell junctions were ablated in the focal plane along a 2 µm line at the centre of adherens junctions for 80 ms at 100pc laser power (10 iterations, thickness 1). The junctions were visualised using cadherin-GFP labelling and were positioned at the centre of the field of view to ensure reproducibility.

Live imaging was performed using a wide-field microscope equipped with a cooled CCD camera (HQ2, Roper Scientific; 64.5 nm/pixel), a GFP filter and HBO illumination to limit photobleaching. Image acquisition was controlled by Metamorph software coupled to ILAS, with images captured every 1.5 s for 7.5 s before ablation, and then again for 31.5 s after ablation.

Post-ablation vertex displacement was quantified over time using the MTrackJ plugin in ImageJ, as previously described (Liang et al., 2016). Data were plotted using Plot Huygens.

#### 9.20 Samples preparation

**Experimental Animals** The animal model used here is Drosophila melanogaster in the context of in vivo/ex vivo experiments. To respect ethical principles, adult flies were anesthetized with CO before any manipulation. To prevent the release of flies outside the laboratory, dead flies were frozen prior to disposal. Stocks of living flies were stored in incubators at either 18 or 25 degrees to maintain optimal conditions for the flies. The fly food contains water, agar (0.8%), sugar (4%), flour (7.4%), yeast (2.8%), mold inhibitor (1%), and propionic acid (0.3%). Genotypes and developmental stages are indicated below. Experiments were performed on both males and females.

**Ex vivo culture of leg imaginal discs**

Leg imaginal discs were dissected at white pupal stage in Schneider's insect medium (Sigma-Aldrich) supplemented with 10% fetal calf serum and 0.6% penicillin-streptomycin as well as 20-hydroxyecdysone at 2 mg/mL (Sigma-Aldrich, H5142). Leg discs were cultured on a slide in 12 L of this medium in a well formed by a 120 mm-deep double-sided adhesive spacer (Secure-SealTM from Sigma-Aldrich) closed with a coverslip. Halocarbon oil was added on the sides of the spacer to prevent dehydration.

**Live imaging and measurements for Drosophila Embryos**

Fly stocks were maintained in incubators at 25°C to ensure optimal developmental conditions. The fluorescent reporter utilized was a w,sqh KI [Tag-RFp-T] 3B line [39]. Embryos were collected 2-4 hours after egg laying (AEL), then dechorionated with bleach diluted by half for 2 minutes.Subsequently, they were mounted between a coverslip and film (Lumox Film 25 ,ref. 94.6077.317, from Sarstedt) with a 120 µm deep spacer (Secure-SealTM from Sigma-Aldrich) in Halocarbon oil.Embryos were oriented and glued on their dorsal side on the film (glue made by incubating Scotch double-sided adhesive tape overnight in heptane).

**Live imaging and measurements on Drosophila pupae**. Cell junctions were visualised thanks to the DE- Cadherin::GFP knock-in. Pupae were picked at white pupae stage and staged for the appropriate time at 25°C. Pupae were then transferred on a piece of double-faced tape onto a microscope slide, dorsal side up. The dorsal part of the pupal case was removed from the head to the most anterior part of the abdomen, while keeping the animal intact. Two spacers, consisting in piles of 4 and 5 18x18 coverslips, were created respectively on the anterior and posterior sides of the pupae. A 24x40 coverslip, on which a thin layer of halocarbon oil had been spread was put onto the two spacers and maintained with nail varnish. The oil should just touch the pupae.
